# Downregulation of *WT1 transcription factor* gene expression is required to promote myocardial fate

**DOI:** 10.1101/2021.07.06.451274

**Authors:** Ines J. Marques, Alexander Ernst, Prateek Arora, Andrej Vianin, Tanja Hetke, Andrés Sanz-Morejón, Uta Naumann, Adolfo Odriozola, Xavier Langa, Laura Andrés-Delgado, David Haberthür, Benoît Zuber, Carlos Torroja, Ruslan Hlushchuk, Marco Osterwalder, Filipa Simões, Christoph Englert, Nadia Mercader

## Abstract

During cardiac development, cells from the precardiac mesoderm fuse to form the primordial heart tube, which then grows by addition of further progenitors to the venous and arterial poles. In the zebrafish, *wilms tumor 1 transcription factor a* (*wt1a*) and *b* (*wt1b*) are expressed in the pericardial mesoderm at the venous pole of the forming heart tube. The pericardial mesoderm forms a single layered mesothelial sheet that contributes to further the growth of the myocardium, and forms the proepicardium. Proepicardial cells are subsequently transferred to the myocardial surface and give rise to the epicardium, the outer layer covering the myocardium in the adult heart. *wt1a/b* expression is downregulated during the transition from pericardium to myocardium, but remains high in proepicardial cells. Here we show that sustained *wt1* expression impaired cardiomyocyte maturation including sarcomere assembly, ultimately affecting heart morphology and cardiac function. ATAC-seq data analysis of cardiomyocytes overexpressing wt1 revealed that chromatin regions associated with myocardial differentiation genes remain closed upon *wt1b* overexpression in cardiomyocytes, suggesting that wt1 represses a myocardial differentiation program. Indeed, a subset of *wt1a/b*-expressing cardiomyocytes changed their cell adhesion properties, delaminated from the myocardial epithelium, and upregulated the expression of epicardial genes, as confirmed by *in vivo* imaging. Thus, we conclude that wt1 acts as a break for cardiomyocyte differentiation by repressing chromatin opening at specific genomic loci and that sustained ectopic expression of *wt1* in cardiomyocytes can lead to their transformation into epicardial cells.

## INTRODUCTION

The heart is one of the first organs to acquire its function and it starts beating long before cardiac development is completed. In mammals, its function is essential to promote blood flow in order to sustain oxygenation and nutrition of the organism. Indeed, heart defects are among the major congenital anomalies responsible for neonatal mortality (1, 2).

The zebrafish is a well-established vertebrate model organism in cardiovascular research given its transparency during early developmental stages and rapid embryonic development (3). Cardiac precursor cells derive from the anterior lateral plate mesoderm (4). At 14 hours postfertilization (hpf), cardiac precursor cells start to express *myosin light chain 7* (*myl7*) (5) and sarcomere assembly begins soon after (6, 7). As the assembly of sarcomeres continues, the cardiac precursor cells migrate and fuse into a cone that later forms the heart tube, which is contractile at 24 hpf and is comprised of a monolayer of cardiomyocytes lined in the interior with an endocardial layer facing the lumen. Next, the heart tube starts to loop, leading to the formation of the two chambers, the atrium and the ventricle (4). Concomitantly, more progenitors enter the heart tube through the arterial and venous poles (8). Around 55 hpf, the outermost cell layer of the heart, the epicardium, starts to form. Epicardial cells arise from the proepicardium, a cell cluster derived from the dorsal pericardium that lies close to the venous pole of the heart. Cells from this cluster are later released into the pericardial cavity and attach to the myocardial surface, forming the epicardium (9, 10).

*Wilms tumor 1 (WT1)* is one of the main epicardial and proepicardial marker genes and plays a central role in epicardium morphogenesis (10, 11). Wt1 contains 4 DNA binding zinc-finger domains in the C-terminus and has been shown to act as a transcription factor (12). *Wt1* is expressed in the epicardium during embryonic development and, in the adult heart, is reactivated after cardiac injury (13).

The zebrafish has two *Wt1* orthologues, *wt1a* and *wt1b* (14). These genes are also expressed in the proepicardium and epicardium (9, 15) in partially overlapping expression domains. Using transgenic reporter and enhancer trap lines (16, 17), we previously showed that *wt1a* and *wt1b* are initially expressed in a few proepicardial cells and later in epicardial cells (18, 19). While *wt1a* and *wt1b* mRNA expression was not detected in the myocardium, *wt1b:eGFP* signal was transiently detected in cardiomyocytes of the atrium close to the inflow tract of the heart. Furthermore, some *wt1a-associated* regulatory regions were found to drive eGFP expression in cardiomyocytes (18). Given that *wt1b* and *wt1a* regulatory elements drive gene expression in the myocardium but endogenous mRNA expression is observed only in the proepicardium and epicardium we hypothesized that *wt1* expression in the myocardium needs to be actively repressed to enable progression of normal heart development needs to be repressed in the myocardium for correct heart development.

To explore whether there is a requirement for *Wt1* downregulation in the myocardium for proper embryonic development, we generated transgenic zebrafish models for tissue specific overexpression of *wt1b* or *wt1a* in cardiomyocytes. We found that sustained *wt1a or wt1b* overexpression in the myocardium induced the delamination and a phenotypic change from cardiomyocytes to epicardial-like cells. Moreover, we observed impaired cardiac morphogenesis, altered sarcomere assembly and delayed myocardial differentiation, which ultimately led to alterations in cardiac function, atrial hypertrophy and fibrosis.

ATAC-seq data analysis of cardiomyocytes overexpressing wt1 revealed that this regulator acts as a break for cardiomyocyte differentiation by reducing chromatin accessibility of genomic loci associated with key processes like sarcomere assembly, establishment of apicobasal polarity and adherens junctions formation.

Altogether, our results demonstrate that transcriptional downregulation of *wt1a/b* expression in cardiomyocytes is a prerequisite for cardiomyocyte specification ensuring correct development of the heart and preventing a phenotypic switch of cardiomyocytes into epicardial cells.

## RESULTS

### Wt1 is downregulated in cardiac progenitors upon their entry into the heart tube

During heart tube growth, cells from the pericardial mesoderm enter the heart tube at the venous pole (8). These cells can be labelled with the line *epi:eGFP* (9), an enhancer trap line of *wt1a* (Fig 1A). We found that during this process, cardiomyocyte precursors downregulate eGFP expression concomitant with the activation of *myl7:mRFP* (Fig 1B and S1 Video). We measured the eGFP/mRFP signal intensity ratio in cells of the hearts tube, from the sinus venosus (SV) towards the growing heart tube. We found that the further away the cells were from the SV, the lower was the eGFP/mRFP ratio (Fig 1C, n=3). To further confirm these observations, we performed SMARTer RNA-seq of cells collected from three distinct regions: pericardium, proepicardium and heart tube at 60 hpf (Fig 1D-F). We detected a gradual decrease in *wt1a* and *wt1b* normalized counts among these three tissues, with the highest counts in PE cells and lowest in cells from the heart tube. The opposite trend was observed for *myl7* expression, being highest in heart tube and lowest in the proepicardium samples.

**Fig 1.**
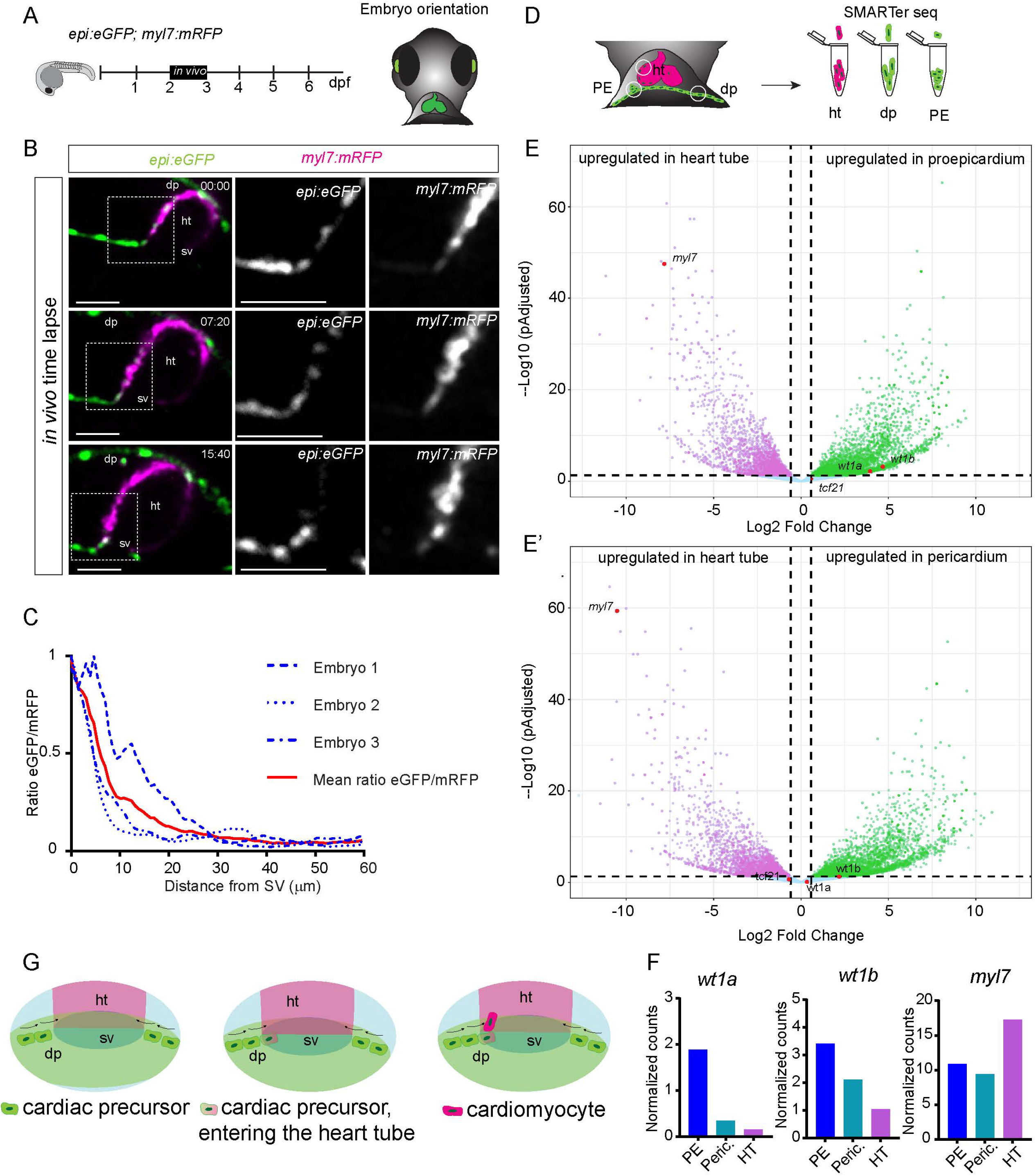
*wt1a* positive cells loose eGFP expression upon entering the heart tube. (A) Schematic representation of the *in vivo* imaging of the developing heart tube. (B) Time-lapse images of the developing heart tube between 52 and 68 hours post fertilization in the double transgenic line *epi:eGFP;myl7:mRFP*. Grey images are single channel zoomed images from the boxes in the merged panels. There is an opposite gradual shift in the expression levels between eGFP and RFP along the time. (C) Quantification of the ratio of eGFP and mRFP levels in cells of the heart tube according to the distance to the sinus venosus (SV). (D) Schematic representation of tissue dissection for SMARTer-seq of pericardium, proepicardium and heart tubes of zebrafish embryos. (E-E’) Volcano plots. Magenta dots indicate upregulated genes in the heart tube. Green dots mark genes upregulated in proepicardium (E) or pericardium (E’). (F) Quantification of normalized counts for the epicardial marker genes *wt1a* and *wt1b*, and the myocardial gene *myl7*. (G) Schematic representation of the downregulation of eGFP and upregulation of mRFP in cardiomyocyte progenitors upon their entry into the heart tube. Scale bars: 50 µm. dp, dorsal pericardium; ht, heart tube; sv, sinus venosus.

We next assessed if the observed downregulation of *Wt1* during cardiomyocyte differentiation from a pool of cardiac precursor cells could be a conserved mechanism during vertebrate heart development and whether this downregulation might be associated with directed repression of *Wt1* gene expression. For this, we inspected previously published data on activating and repressing Histone marks at the Wt1 genomic locus during four stages of cardiac differentiation from mouse embryonic stem cells (mESCs) (20) (S1 Fig). Low levels of *Wt1* transcripts were visible in mESCs but absent throughout differentiation stages. Similarly, Histone 3 K27 acetylation (H3K27ac) enrichment - which correlates with active promoter and enhancer activity – was present in regions proximal to the *Wt1* transcriptional start site (TSS) in mESCs and mesodermal progenitor cells. Interestingly, some of these regions also co-localized with enhancer elements known to drive epicardial-specific reporter gene expression (21). Conversely, Histone H3 K27 trimethylation (H3K27me3) signatures, which associate with repressed regions, were near-absent in mESCs, while in cardiac precursors cells and differentiated cardiomyocytes, they massively decorated the extended regions flanking the *Wt1* TSS, including the epicardial enhancer elements. Together, these observations indicate that during cardiomyocyte differentiation, *Wt1* expression and epicardial enhancers become actively repressed.

In summary, during heart development, cardiac precursor cells downregulate *wt1* upon their entry into the heart tube, which might be a prerequisite for their differentiation into cardiomyocytes (Fig 1G).

### Cardiomyocytes that overexpress *wt1* can delaminate from the heart, are depleted of sarcomeric proteins and start expressing epicardial markers

We next aimed at exploring the biological relevance for the observed downregulation of *wt1* in cardiomyocytes. Therefore, to analyze the consequence of *wt1* expression in cardiomyocytes, we generated the line *Tg(b-actin2:loxP-DsRED-loxP-eGFP-T2A-wt1a)*, to conditionally induce the expression of *wt1a.* Crossing this line into *Tg(myl7:CreERT2)* (22) allowed the temporally induced overexpression of *wt1a* in cardiomyocytes. Hereafter, the double transgenic line is called *myl7:CreERT2;eGFP-T2A-wt1a*. We administered 4-hydroxytamoxifen (4-OHT) from 24 hpf to 4 days postfertilization (dpf) to induce recombination of loxP sites and activation of *wt1a* and *eGFP* expression during embryogenesis in cardiomyocytes (S2 Fig). We confirmed *wt1a* and *eGFP* overexpression in the heart by RT-qPCR (S2 Fig). Comparison of *eGFP* and *wt1a* expression between *myl7:CreERT2;eGFP-T2A-wt1a* with and without 4-OHT administration revealed a 4-fold increase in *eGFP* and *wt1a* expression in the latter (S2 Fig). Moreover, we generated a line to overexpress *wt1b*. We decided to use the Gal4/UAS system in this case, to allow a more homogeneous expression in the myocardium. The line *Tg(eGFP:UAS:wt1b)* allowed overexpression of *wt1b* and *eGFP* under a bidirectional *UAS* promoter. We crossed the line into *Tg(myl7:Gal4)* (23); the double transgenic line will be hereafter called *myl7:Gal4;eGFP:UAS:wt1b* (S3 Fig). *wt1b* and *eGFP* expression in cardiomyocytes of the double transgenic line *myl7:Gal4;eGFP:UAS:wt1b* was four-fold upregulated compared to cells from the single transgenic eGFP:UAS:*wt1b* (S2 Fig). As a control, we used the double transgenic line *Tg(eGFP:UAS:RFP);(myl7:Gal4)* (24), hereafter named *myl7:Gal4;eGFP:UAS:RFP*. RT-qPCR analysis also indicated that expression of *wt1b* and *wt1a* could be monitored via GFP imaging (S2 Fig).

Using these new lines, we analyzed the effect of sustained *wt1a* and *wt1b* overexpression in cardiomyocytes during heart development (Fig 2A). In *wt1a-*overexpressing hearts but not in controls, we were able to observe eGFP-positive cardiomyocytes located at an apical position, protruding towards the pericardial cavity at 5 dpf (Fig 2B-C’’’). Moreover, these delaminating cardiomyocytes showed reduced expression of Myosin Heavy Chain (MHC), suggesting that they lost to some extent a myocardial phenotype (Fig 2C-C’’’). We quantified how many of the delaminating cells were GFP^+^ or GFP^-^ and found that only GFP^+^ cells were delaminating, indicating that this delamination process is due to a cell-autonomous effect of *wt1a* in cardiomyocytes (S1 Table). We detected a similar occurrence in *wt1b* overexpressing hearts, starting at 3 dpf (Fig 2D-E’’’). Some GFP-positive cells delaminated towards the apical myocardial surface. Those cells were not positive for the myocardial marker MHC.

**Fig 2.**
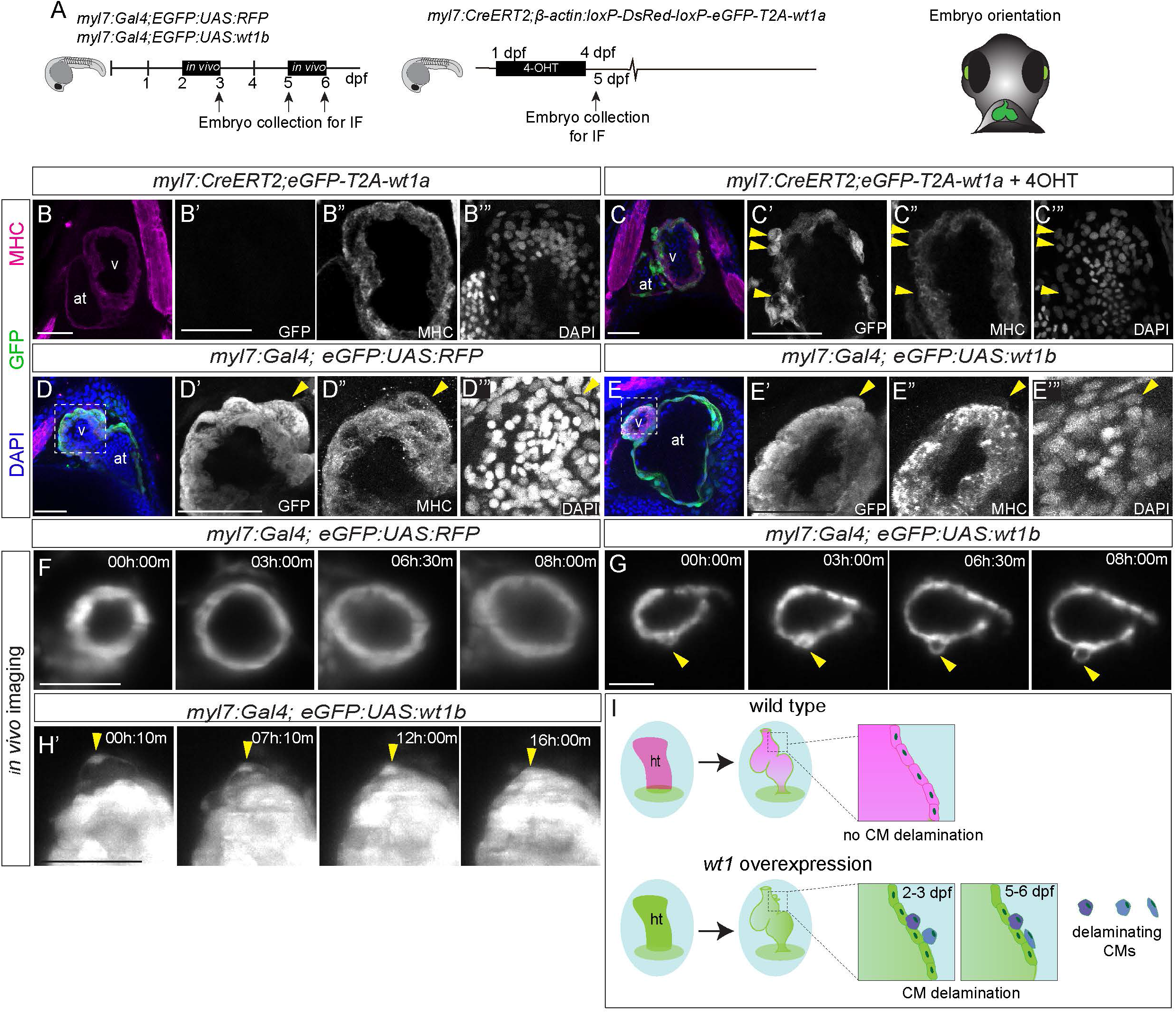
Ventricular cardiomyocytes delaminate from the ventricle and change their fate upon *wt1b* overexpression. (A) Schematic representation of used transgenic lines and position of the embryos for imaging. (B-C’”) Whole mount immunofluorescence against GFP (green) and MHC (magenta) on *myl7:CreERT2,eGFP-T2A-wt1a* hearts at 5 days post fertilization (dpf), non-recombined (B-B’’’) and recombined by addition of 4-OHT between 24 hours post fertilization (hpf) and 4 dpf (C-C’’’). Shown are maximum intensity projections of 5 optical sections with a distance of 1.5 µm between two consecutive sections. Yellow arrows point GFP positive cardiomyocytes located on the apical myocardial surface revealing reduced MHC staining. (D-E’’’) Whole mount immunofluorescence against GFP and MHC on a *myl7:Gal4;eGFP:UAS:RFP* (D-D’’’) and a *myl7:Gal4;eGFP:UAS:wt1b* (E-E’’’) embryo, at 3 dpf. DAPI was used for nuclear counterstain. Shown are maximum intensity projections of 20 stacks with a distance of 1 µm between two consecutive optical sections of the heart region. (D-D’”) and (E-E’”) are magnifications of the area of the ventricle marked by the dashed bounding boxes in D and E, respectively. Yellow arrowhead points to a GFP-positive cell that is MHC^+^ in D-D’’’ and to a GFP+/MHC-cell in E-E’’’). (F-G) Time lapse images of the ventricle in a *myl7:Gal4; eGFP:UAS:RFP* (F) or *myl7:Gal4; eGFP:UAS:wt1b* (G) embryo between 2 and 3 dpf. Elapsed time since initial acquisition is stamped in each panel. Arrowhead in G point to a cell extruding from the ventricle. (H) Time lapse images of the ventricle in a *myl7:Gal4; eGFP:UAS:wt1b* embryo between 5 and 6 dpf. Elapsed time since initial acquisition is stamped in each panel. Note how a delaminating cell changes morphology along time and flattens down (yellow arrowhead). (I) Model of the delamination process of *wt1b* overexpressing cardiomyocytes. Scale bar, 50 µm. at, atrium; CM, cardiomyocyte; ht, heart tube; IF, immunofluorescence; v, ventricle

To better understand the origin of these apically positioned eGFP-positive cells, in the *wt1b* overexpression hearts, we performed *in vivo* imaging in *myl7:Gal4;eGFP:UAS:RFP* and *myl7:Gal4;eGFP:UAS:wt1b* between 2 and 3 dpf (Fig 2F, G and S2 Video). In *myl7:Gal4:eGFP:UAS:wt1b* hearts, some eGFP-positive cells started to round up and initiated delamination from the myocardium. Cells gradually changed from a flat to a rounded shape and ultimately remained adherent to the outer myocardial layer (Fig 2G and S2 Video; n=4). This event of cell delamination was not observed in *myl7:Gal4:eGFP:UAS:RFP* control embryos (Fig 2F and S2 Video; n=2). Apical extrusion of cardiomyocytes can be a consequence of myocardial malformation during which extruded cells are eliminated (25, 26). However, here we found that the delaminated cells did remain attached to the myocardial surface. Between 5 and 6 dpf, these delaminated cells lost their rounded shape and flattened, acquiring an epicardial-like morphology (Fig 2H-I and S3 Video 3, n=4).

To confirm that this type of cellular delamination with loss of MHC expression was specific of the overexpression of *wt1* we generated the *Tg(eGFP:UAS:tcf21),* which we then crossed into the *Tg(myl7:Gal4)* (S3 Fig). This double transgenic allowed us to overexpress another well-known epicardial marker, *tcf21* (27) in cardiomyocytes. Contrary to what we observed when overexpressing *wt1a/b* in cardiomyocytes (S3 Fig), in the large majority of these embryos (59/62) we did in not observe apical delamination in the hearts of the *myl7:Gal4;eGFP:UAS:tcf21* fish. In the very few cases where delamination occurred (3/62), the protruding cells still expressed MHC (S3 Fig).

Due to the position and change of morphology of delaminated cells we hypothesized, these cells had undergone a change of fate. For a better characterization of a possible switch to an epicardial fate we performed immunofluorescence labeling with the epicardial markers Aldehyde dehydrogenase 2 (Aldh1a2) (28, 29) and Caveolin 1 (Cav1) (30) (Fig 3A). We detected GFP/Aldh1a2 double positive cells in *wt1b* overexpression hearts (n=3) but not in controls (n=4) (Fig 3B-C’’’). Similarly, whereas in control hearts (n=6) we could not observe GFP/Cav1 double positive cells (Fig 3D-D’’’), in *wt1b*-overexpressing hearts (n=4), we identified GFP positive cells that also expressed Caveolin 1 (Fig 3E-E’’’). We also detected eGFP^+^ cells within the epicardium of *wt1a* overexpressing hearts, but not in controls (Fig 3F-G, H-H’, I-J’, K-K’, L-M’, N-N’, O-P’ and Q-Q’). These eGFP^+^ cells did not express MHC (Fig 3J”, K”, P” and Q”), and were Aldh1a2-positive (Fig 3J’’’ and L’’’) as well as Caveolin 1 positive (Fig 3P’’’ and Q’’’) strongly suggesting that *wt1a* overexpressing cardiomyocytes switched their fate to epicardial cells. We tested for the colocalization of *wt1a*-expression in cardiomyocytes with a third epicardial marker, *transglutaminase b* (*tgm2b)* (31). We performed *in situ* hybridization against *tgm2b* mRNA followed by immunohistochemistry against eGFP (S4 Fig). In non-recombined *myl7:CreERT2; eGFP-T2A-wt1a* hearts, *tgm2b* expression was only visible in few epicardial cells in ventricle and we could not observe any co-localization with eGFP expressing cells (S4 Fig). However, in embryonically recombined *myl7:CreERT2; eGFP-T2A-wt1a* hearts, we could observe cells co-expressing *tgm2b* and eGFP located within the epicardium (S4 Fig).

**Fig 3.**
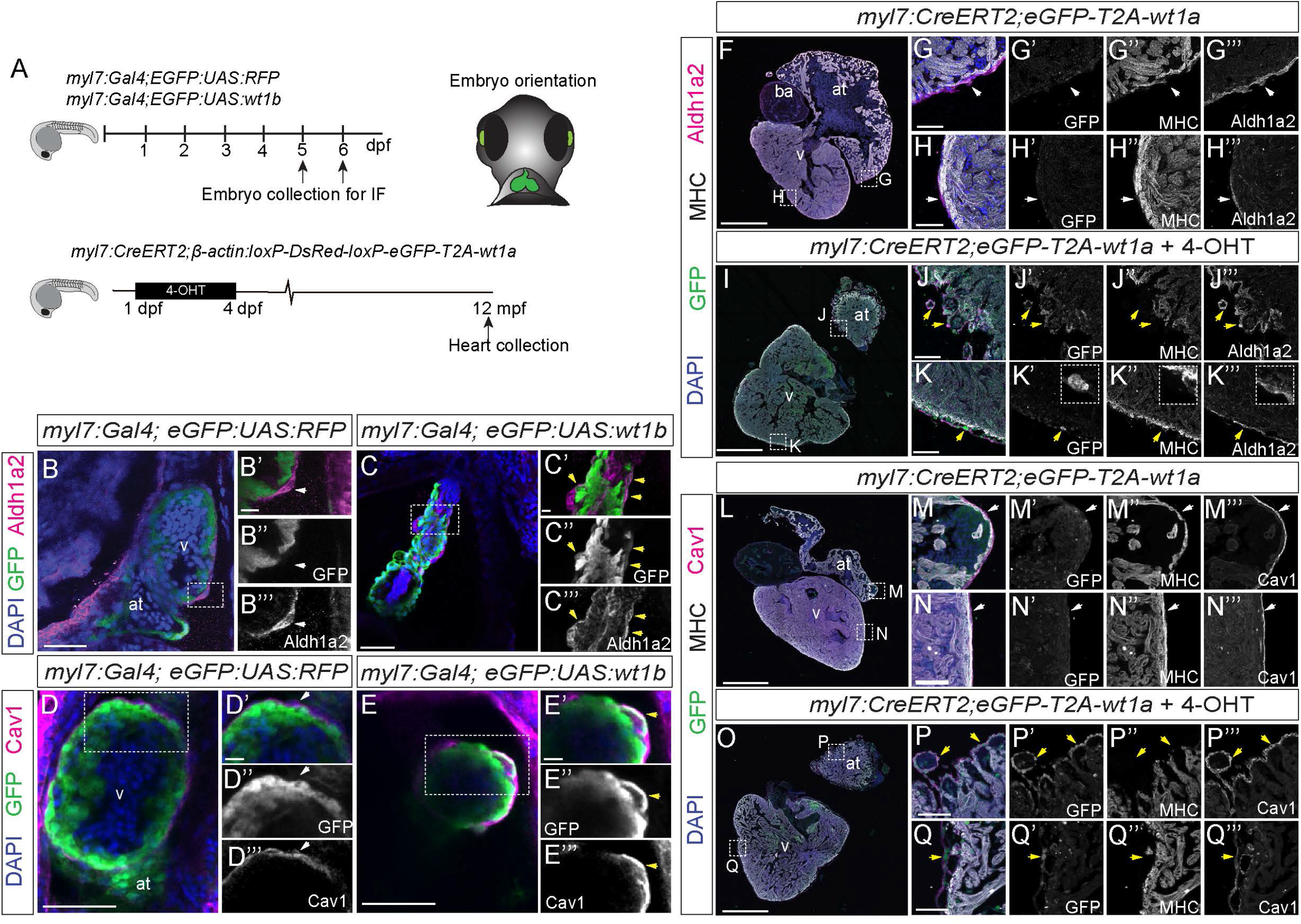
Delaminated *wt1* overexpressing in cardiomyocytes start to express epicardial markers. (A) Schematic representation of the lines used and the time points during which 4-hydroxytamoxifen (4-OHT) was administered to *myl7:CreERT2;β-actin:loxP-DsRed-loxP-eGFP-T2A-wt1a* fish (in short *myl7:CreERT2,eGFP-T2A-wt1a*), as well as embryo orientation for image acquisition. (B-C’’’) Whole mount immunofluorescence against GFP and Aldh1a2 in a *myl7:Gal4;eGFP:UAS:RFP* (B-B’’’) and *myl7:Gal4;eGFP:UAS:wt1b* (C-C’’’) embryo, at 5 dpf. Shown are maximum intensity projections of 5 images with a distance of 1 µm between two consecutive optical sections. (B’-B’’’) Zoomed view of the boxed area in B. (C’-C’’’) Zoomed view of the boxed area in C. White arrow, epicardial cells positive for Aldh1a2 and negative for GFP. Yellow arrows, epicardial cells that express both Aldh1a2 and GFP. Green, GFP; magenta, Aldh1a2; blue, DAPI. (D-E’’’) Whole mount immunofluorescence against GFP and Caveolin 1 (Cav1) in a *myl7:Gal4;eGFP:UAS:RFP* (D-D’’’) and *myl7:Gal4;eGFP:UAS:wt1b* (E-E’’’) embryo, at 6 dpf. Shown are maximum intensity projections of 10 consecutive optical section with a distance of 1.5 µm between them. (D’-D’’’) Zoomed view of the boxed area in D. (E’-E’’’) Zoomed view of the boxed area in E. White arrows, epicardial cells positive for Cav1 and negative for GFP. Yellow arrows, epicardial cells that express both Cav1 and GFP. Green, GFP; magenta, Cav1; blue, DAPI. (F-K’’’) Immunofluorescence against GFP (green), MHC (white) and Aldh1a2 (magenta) on paraffin sections of (F-H’’’) *myl7:CreERT2;eGFP-T2A-wt1a* and (I-K’’’) *myl7:CreERT2;eGFP-T2A-wt1a* + 4-OHT adult hearts. Shown are sections of the heart (F and I), as well as zoomed views of indicated regions (G-G’’’, H-H’’’, J-J’’’ and K-K’’’). Both, merged and single channels are shown, as indicated in the panel. White arrowheads, cells positive for Aldh1a2 only. Yellow arrowheads point to cells positive for GFP and Aldh1a2 signal that lack MHC staining, and which are located close to the myocardial surface. (L-Q’”) Immunofluorescence against GFP (green), MHC (white) and Caveolin 1 (Cav1) (magenta) on paraffin sections of (L-N’’’) *myl7:CreERT2;eGFP-T2A-wt1a* and (O-Q’’’) *myl7::CreERT2;eGFP-T2A-wt1a* + 4-OHT adult hearts. Shown are sections of the heart (L and O), as well as zoomed views of indicated regions (M-N’’’ and P-Q’’’). Both, merged and single channels are shown, as indicated in the panel. White arrowheads point to cells positive only for Cav1. Yellow arrowheads point to cells positive for GFP and Cav1 signal that lack MHC staining, and which are located close to the myocardial surface. Scale bars: 500 µm (F, I, L, O) 50 µm (B, C, D, E, G, H, J, K, M, N, P, Q) and 10 µm (B’-B”’, C’-C’’’, D’-D”’ and E’-E’’’). at, atrium; Cav1, Caveolin1; MHC, Myosin Heavy Chain; v, ventricle.

These results suggest that upon sustained ectopic overexpression of *wt1a/b,* cardiomyocytes can delaminate apically from the myocardial layer and adopt features of epicardial cells that contribute to the formation of the epicardial layer even in the adult heart.

### *wt1b* overexpression disrupts cell-cell contacts and the basement membrane of the cardiomyocytes

We decided to get a better understanding on the cellular mechanisms underlying cardiomyocyte apical delamination of cardiomyocyte upon *wt1* expression (Fig 4A). Previous reports showed that correct development and morphogenesis of the heart requires cell-cell adhesion and polarization of the cardiomyocytes (32). The proper localization of tight junctions and adherens junctions has conventionally been used to assess the polarization of the cells (33). We first performed immunostainings against ZO-1, a component of tight junctions (34) (Fig 4B-E’). Whereas the *myl7:Gal4:eGFP:UAS:RFP* control hearts (n=6) showed discrete apical localization of ZO-1 (Fig 4B-C’), in *myl7:Gal4:eGFP:UAS:wt1b* hearts (n=8) ZO-1 levels were reduced, the signal was diffuse and not clearly localized to apical junctions between cardiomyocytes (Fig 4D-E’). This suggests defects in the formation and localization of tight junctions upon *wt1b* overexpression. To evaluate the formation of adherens junctions, we crossed the *Tg(myl7:cdh2-tdTomato)^bns78^*line (35) with *myl7:Gal4:eGFP:UAS:wt1b*. This allowed us to specifically visualize subcellular localization of cdh2-tdTomato in *wt1b*-overexpressing cardiomyocytes and control siblings (Fig 4A and F-M’). At 5 dpf, in control embryos (n=6), tdTomato signal was clearly localized to cell-cell junctions (Fig 4F-G) and detected apically in cardiomyocytes (Fig 4H-I’). In contrast, *myl7:cdh2-tdTomato;myl7:Gal4:eGFP:UAS:wt1b* hearts (n=8), showed a diffused and patchy staining for cdh2-tdTomato, which was not restricted to the apical side of the cardiomyocytes (Fig 4J-J’’ and L-M’). Moreover, we observed loss of cdh2-tdTomato signal in the delaminating cells further indicating a loss of polarity in these extruding cells (Fig 4K-K”). To confirm the impairment in the formation of adherens junctions we did an immunostaining against β-catenin, a core component of adherens junctions (36). Similar to what we had observed for cdh2-td-Tomato, β-catenin staining was located at the apical side of cardiomyocytes in *myl7:Gal4:eGFP:UAS:RFP* control hearts (n=5) (Fig 4N-O’). However, in *wt1b*-overexpressing hearts (n=5) β-catenin staining was no longer detected (Fig 4P-Q’). Taken together, this data shows that sustained expression of *wt1b* in cardiomyocytes leads to the mislocalization of tight junctions and adherens junctions, indicating an impairment of the apical domain in cardiomyocytes.

**Fig 4.**
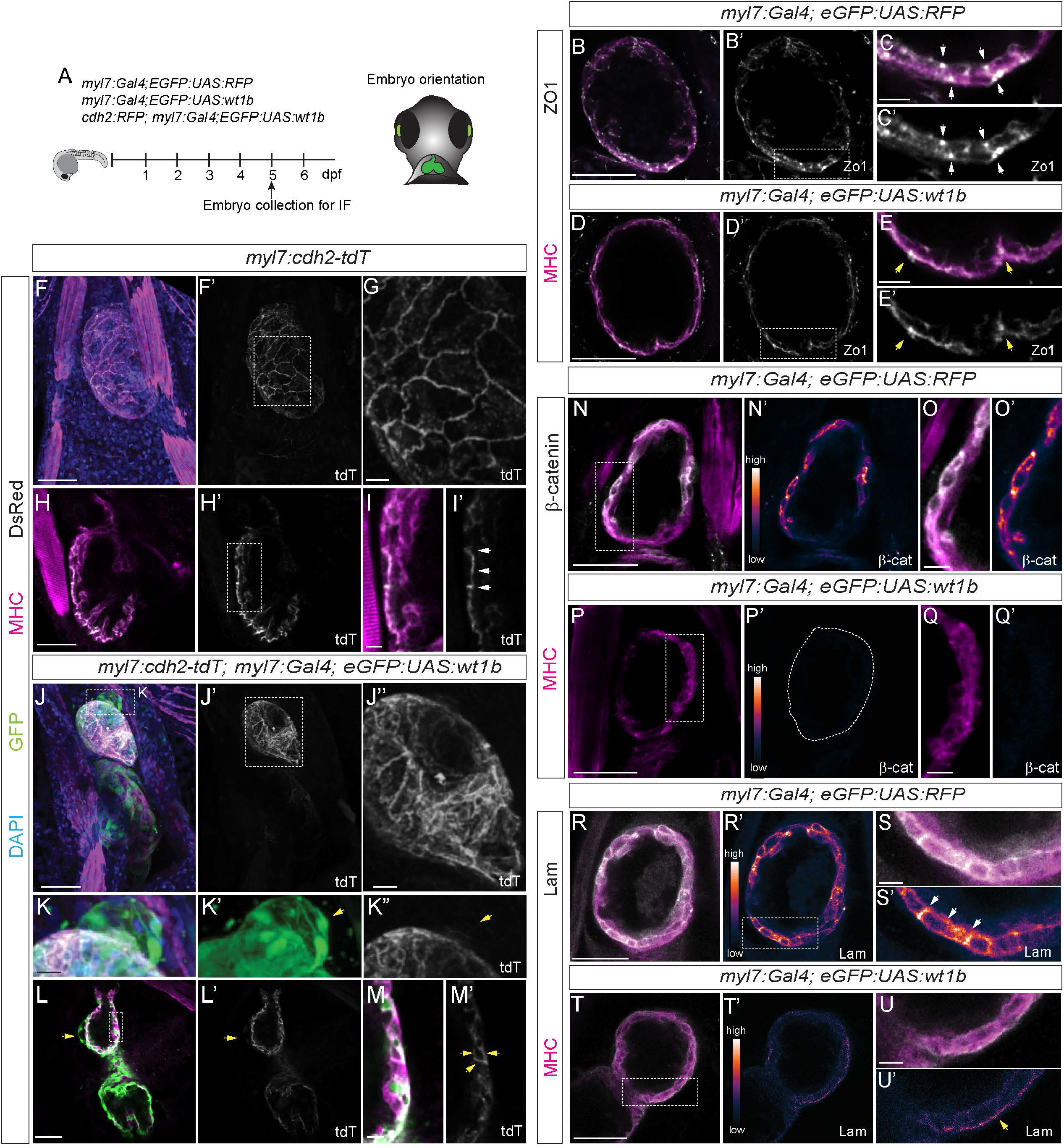
Expression of cell junction and polarity markers in *wt1b*-overexpressing hearts. (A) Schematic representation of the lines used and embryo orientation for imaging. (B-E’) Immunofluorescence against zonula occludens 1 (ZO-1) and myosin heavy chain (MHC) in 5 days post fertilization (dpf) *myl7:Gal4;eGFP:UAS:RFP* (B-C’) and *myl7:Gal4;eGFP:UAS:wt1b* (D-E’) embryos. Shown are sagittal single planes of the ventricle. Single channels (B’, C’, D’ and E’) show ZO1 signal. (C-C’) Zoomed views of the box in B’. White arrows point to ZO1 signal. (E-E’) Zoomed views of the box in D’. Yellow arrows point to ZO1 signal. (F-M’) Immunofluorescence against tdTomato (tdT) (using a DsRed antibody) and MHC in 5 dpf *myl7:cdh2-tdTomato* (F-I’) and *myl7:cdh2-tdTomato*;*myl7:Gal4;eGFP:UAS:wt1b* (J-M’) embryos. (F-G) 3D projections of a heart. (G) Zoomed view of the box region in F’. (H-I’) Sagittal single planes of the ventricle. (I-I’) Zoomed view of the box in H’. White arrows point do regions with tdT signal. (J-K”) 3D projections of a heart. (J”) Zoomed view of the box in J’. (K-K”) Zoomed views of the box in J. Yellow arrows point to delaminating cells from the ventricle. Note the absent tdT signal from the delaminated cells (K”). (L-M’) Sagittal single planes of the ventricle. (M-M’) Zoomed view of the box in L. Yellow arrows highlight tdT signal. (N-Q’) Immunofluorescence against Beta-catenin (β-cat) and myosin heavy chain (MHC) in 5 dpf *myl7:Gal4;eGFP:UAS:RFP* (N-O’) and *myl7:Gal4;eGFP:UAS:wt1b* (P-Q’) embryos. Shown are sagittal single planes of the ventricle. Single channels (N’, O’, P’ and Q’) show β-cat signal. LUT color shows gradient of β-cat signal intensity. Lower signal is in blue and the higher signal in orange to white. (O-O’) Zoomed views of the box in N. (Q-Q’) Zoomed views of the box in P. Marked region in P’ indicates the ventricle. (R-U’) Immunofluorescence against Laminin (Lam) and MHC in 5 dpf *myl7:Gal4;eGFP:UAS:RFP* (R-S’) and *myl7:Gal4;eGFP:UAS:wt1b* (T-U’) embryos. Shown are sagittal single planes of the ventricle. Single channels (R’, S’, T’ and U’) show Laminin staining. LUT color shows gradient of laminin signal intensity. Lower intensity is in blue and the higher intensity in orange to white. (S-S’) Zoomed views of the box in R’. White arrows highlight Laminin signal. (U-U’) Zoomed views of the box in P. Yellow arrows highlight Laminin signal. Scale bar: 50 µm (B-B’, D-D’, F-F’, H-H’, J-J’, L-L’, N-N’, P-P’, R-R’ and T-T’); 10 µm (C-C’, E-E’, G, I-I’, J”-K”, M-M’, O-O’, Q-Q’, S-S’ and U-U’).

Lam, Laminin; tdT, tdTomato

To understand the basal domain landscape of cardiomyocytes we did an immunostaining against Laminin, a component of the basement membrane. Laminins have been associated with myocardial differentiation and with regulating the sarcolemmal properties (37–40). At 5 dpf, in the hearts of control fish (n=5) we observed clear anti-Laminin staining at the basal and lateral domains of cardiomyocytes (Fig 4 R-S’), which correlates with previous observations (37, 41). Laminin expression levels were severely reduced in *wt1b* overexpression hearts (n=5), with no Laminin observed in the lateral domains of the cardiomyocytes (Fig 4T-U’). Thus, the observed reduced levels of Laminin and its impaired deposition upon *wt1b* overexpression point towards an improper basal domain of cardiomyocytes.

Taken together, our observations indicate that cardiomyocyte apicobasal polarization may be disrupted upon *wt1b* overexpression.

### Overexpression of *wt1* in cardiomyocytes hinders cell maturation and disrupts its structural organization

The disruptions in cell junctions and cell extrusion that we observed in *wt1b* overexpressing cardiomyocytes led us to question the maturation and general architecture of these cells. Using whole mount immunofluorescence, we observed reduced MHC staining in ––*wt1b-*overexpressing hearts at 1 dpf when compared to controls (Fig 5A-C’). The reduction of MHC staining was specific to the heart, as it was not observed in the skeletal muscle of the myotome (Fig 5D). Although at 6 dpf we observed an increase in the levels of MHC signal in *wt1b* overexpressing cardiomyocytes, the levels never reach those observed in the control group (Fig 5E-G). We also analyzed *myl7* mRNA expression levels using whole mount *in situ* hybridization. Consistent with the results obtained using MHC immunostaining, at 3 dpf, *myl7* expression was reduced in *myl7:Gal4;eGFP:UAS:wt1b* (23/25) compared to their single transgenic *eGFP:UAS:wt1b* control siblings (Fig 5H-I). We reasoned that the reduced levels in MHC and *myl7* staining could be indicative of an impaired maturation of cardiomyocytes. To test this hypothesis, we performed immunofluorescence staining against Alcam, a marker for undifferentiated cardiomyocytes (42, 43). At 6 dpf, we observed higher Alcam staining levels in hearts overexpressing *wt1b* when compared to control hearts (Fig 5J-L).

**Fig 5.**
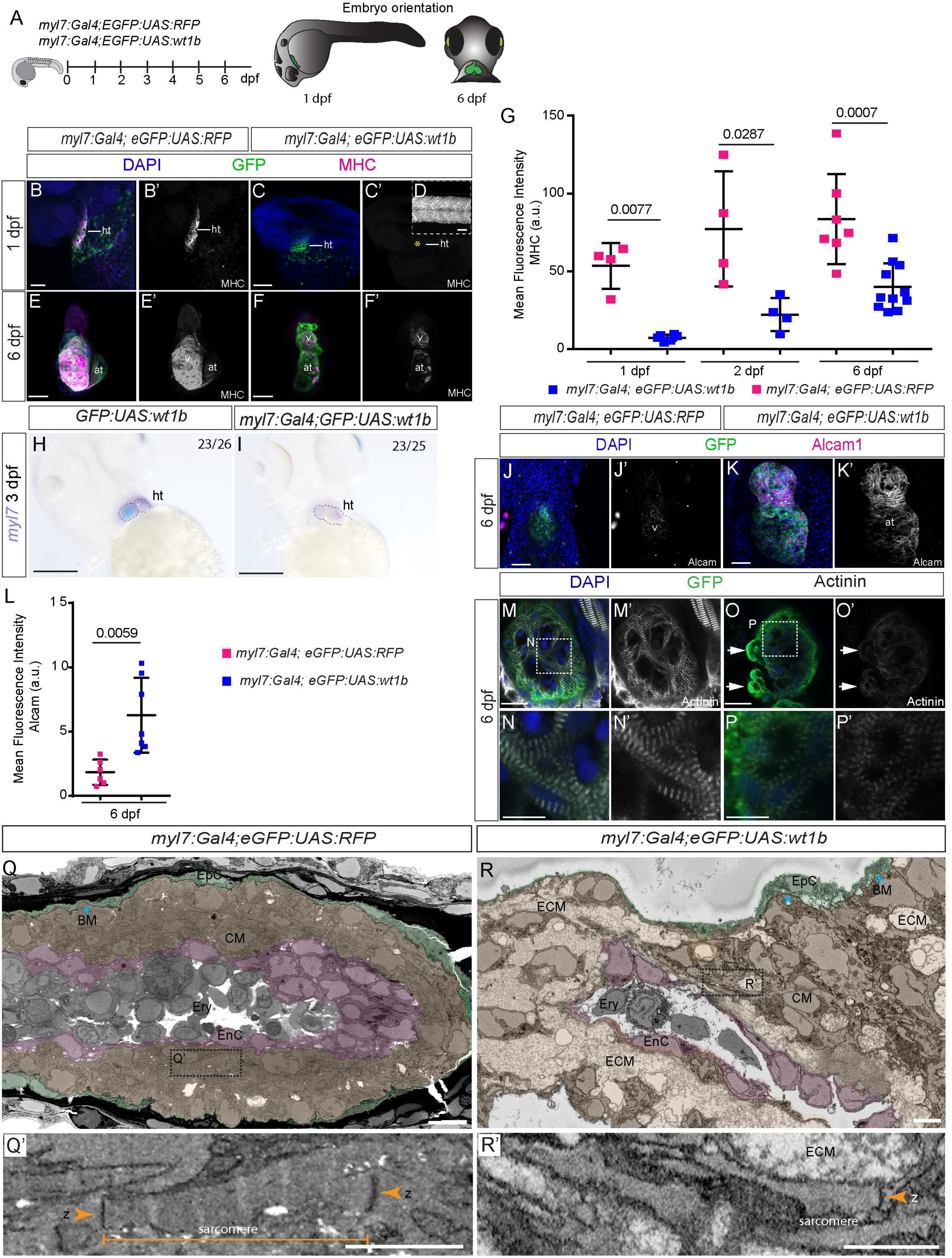
Changes in cardiomyocyte maturation and structure upon *wt1b* overexpression. (A) Schematic representation of the lines used and embryo orientation for stainings and imaging. (B-F’) Immunofluorescence against GFP and myosin heavy chain (MHC) on *myl7:Gal4; eGFP:UAS:RFP* and *myl7:Gal4;eGFP:UAS:wt1b* zebrafish embryos. (B-C’) 3D projection of 1 day post fertilization (dpf) embryos. Shown are lateral views of the cardiac tube. Yellow asterisk in C’ indicates absent MHC staining in the heart. (D) MHC staining of the myotome region of the *myl7:Gal4; eGFP:UAS:wt1b* embryo at 1 hpf. (E-F’) 3D projections of the heart region at 6 dpf. Shown are ventral views, the head is to the top. (G) Quantification of mean fluorescence intensity in the heart region for *myl7:Gal4; eGFP:UAS:RFP* and *myl7:Gal4;eGFP:UAS:wt1b* zebrafish, at indicated developmental stages. Statistical significance was calculated by unpaired t-test, with Welch’s correction (24 hpf) and unpaired t-test for the remaining group comparisons. Means ±SD as well as individual measurements are shown. (H-I) Whole mount mRNA *in situ* hybridization against *myl7* mRNA in (H) *eGFP:UAS:wt1b* and (I) *myl7:Gal4;eGFP:UAS:wt1b* zebrafish embryos at 3 dpf. Embryos are positioned ventrally, with the head to the top. (J-K’) Immunofluorescence against GFP and Alcam on *myl7:Gal4;eGFP:UAS:RFP* and *myl7:Gal4;eGFP:UAS:wt1b* zebrafish embryos. Shown are 3D projection of the heart region in a 6 dpf old larva (ventral views, the head is to the top). (L) Quantification of mean fluorescence intensity of anti-Alcam staining as shown in K-L’. Statistical significance was calculated by unpaired t-test, with Welch’s correction. Shown are mean ±SD as well as individual measurements. (M-P’) Immunofluorescence against GFP and Actinin. Shown are maximum intensity projections of two consecutive optical sections with a step size of 2 µm of the ventricle of *myl7:Gal4;eGFP:UAS:RFP* (M-N’) and *myl7:Gal4;eGFP:UAS:wt1b* (O-P’) at 6 dpf. (N,N’ and P,P’) Maximum intensity projections of boxed regions in (M) and (O), respectively. (Q-R’) Serial block face scanning electron microscope images of zebrafish hearts. Shown are single sections of *myl7:Gal4;eGFP:UAS:RFP* (Q-Q’) and *myl7:Gal4;eGFP:UAS:wt1b* (R-R’) hearts. Different cell layers are highlighted with colors. (Q’ and R’). Zoomed areas highlighting sarcomeres. Green labels the epicardium, magenta marks the endocardial layer and orange highlights the myocardium. Orange arrowheads, z-bands; Cyan arrowhead, basement membrane delimiting epicardium and myocardium. Scale bars, 50 µm (B-F’H-M and O-O’); 1 µm (Q and R), 500 nm (Q’ and R’), 10 µm (N, N and P,P’). at, atrium; BM, basement membrane; CM, cardiomyocyte; dpf, days post fertilization; ECM, extracellular matrix; EnC, endothelial cell; EpC, epicardial cell; Ery, erythrocyte; v, ventricle; ; z, z-line. Green, GFP; magenta, MHC, Alcam; blue, DAPI.

We next analyzed if sarcomere assembly was impaired in *myl7:Gal4;eGFP:UAS:wt1b* animals. We performed immunofluorescence staining against Actinin, a protein known to be produced in the z line of the sarcomeres (44). Qualitative assessment of Actinin revealed that not only the its levels were lower but also the z-lines were thicker and shorter (Fig 5M-P’) upon myocardial *wt1b* overexpression. Z-line disruption was particularly evident in delaminating cardiomyocytes (Fig 5O). We next sought to analyze sarcomere structure more in detail using serial block face scanning electron microscopy (SBFSEM) (Fig 5Q-R’and S4, S5, S6 and S7 Video). Z-bands were present at the sarcomere boundaries in both groups. While sarcomeres could be easily followed from z-band to z-band in the control heart (Fig 5Q-Q’ and S5 Video), this was not possible in *wt1b* overexpressing hearts (Fig 5R-R’and S7 Video). A further ultrastructural defect we observed in the *wt1b* overexpression heart was the presence of large intercellular spaces of extracellular matrix, between cardiomyocytes, the epicardium, and endocardium. Moreover, while the control heart revealed a clearly visible basement membrane between the epicardium and the myocardium as well the endocardium and myocardium (dark black line), this structure was not always visible in the *wt1b-*overexpressing heart (Fig 5A-B’and S3 and S5 Video). This observation correlates with the impairment in Laminin staining reported in *wt1b*-overexpressing cardiomyocytes (Fig 4P-S’).

Altogether, our findings indicate that sustained expression of *wt1b* in cardiomyocytes affects heart development leading to impaired cardiomyocyte maturation and negatively affecting the cardiac ultrastructure, including cardiomyocyte sarcomere assembly and the extracellular matrix.

### Overexpression of wt1b in cardiomyocytes results in global reduced chromatin accessibility

Seeing that *wt1b* overexpression in cardiomyocytes induced several cardiac malformations and caused a phenotypic change in some cells we decided to explore how the sustained expression of this transcription factor was affecting chromatin accessibility. To this purpose, we performed Assay for Transposase-Accessible Chromatin sequencing (ATAC-seq) (45) in 5 dpf, FAC sorted GFP^+^ cells from either the *myl7:Gal4;eGFP:UAS:RFP* control or *myl7:Gal4;eGFP:UAS:wt1b* larvae (Fig 6A). We identified 1452 differential peaks in *wt1b* overexpressing cardiomyocytes, of which almost all except for 14 peaks showed reduced chromatin accessibility (Fig 6B S1 Data). Most of the differential accessible regions were located close to promoter regions (38.87%), in introns (30.37%) or in distal intergenic regions (26.14%) (Fig 6C). We performed Gene Ontology (GO) analysis for the genes lying in close proximity to the differentially accessible regions. From the top 25 Biological Pathways that had reduced accessibility of regions in close proximity of genes associated with these pathways, five of them account for muscle development (Fig 6D and S2 Data). Within the top 25 Cellular Component pathways we found some to be involved in “actin cytoskeleton”, “basolateral plasma membrane”, “apical part of the cells”, “contractile fiber” or “myofibril” (Fig 6E and S2 Data). Within the top 25 Molecular Function pathways (Fig 6F and S2 Data) five of them are directly implicated in transcription regulation and another four in cytoskeleton formation and cell adhesion, such as “actin binding”, “actin-filament binding”, “cell adhesion molecule binding” and “beta-catenin binding”. All of these pathways, which are underrepresented in the *myl7:Gal4;eGFP:UAS:wt1b* samples strongly correlate with the defects observed in hearts overexpressing *wt1b*. To identify potential transcription factors that might be binding to the differentially accessible regions, we performed MEME-Centrimo motif analysis, and found WT1 to be one of the top 5 motifs represented (E-value = 7.8e-5). This motif could be identified in 672 (46.25%) of the differentially accessible regions (3 open and 669 closed) (Fig 6G-G’). To further investigate which of the open regulatory regions and their associated genes were potential direct targets of WT1, we compared our ATAC-seq data with WT1 target genes identified in the CHIP-atlas database (46). 41% of the regions associated with differential accessibility (426, of which only 6 represent regions with open chromatin) identified in our ATAC-seq were shared with the CHIP-atlas database for WT1. GO analysis of the associated common genes identified pathways similar to those observed previously, suggesting a direct regulation of these pathways by Wt1b (S5 Fig).

**Fig 6.**
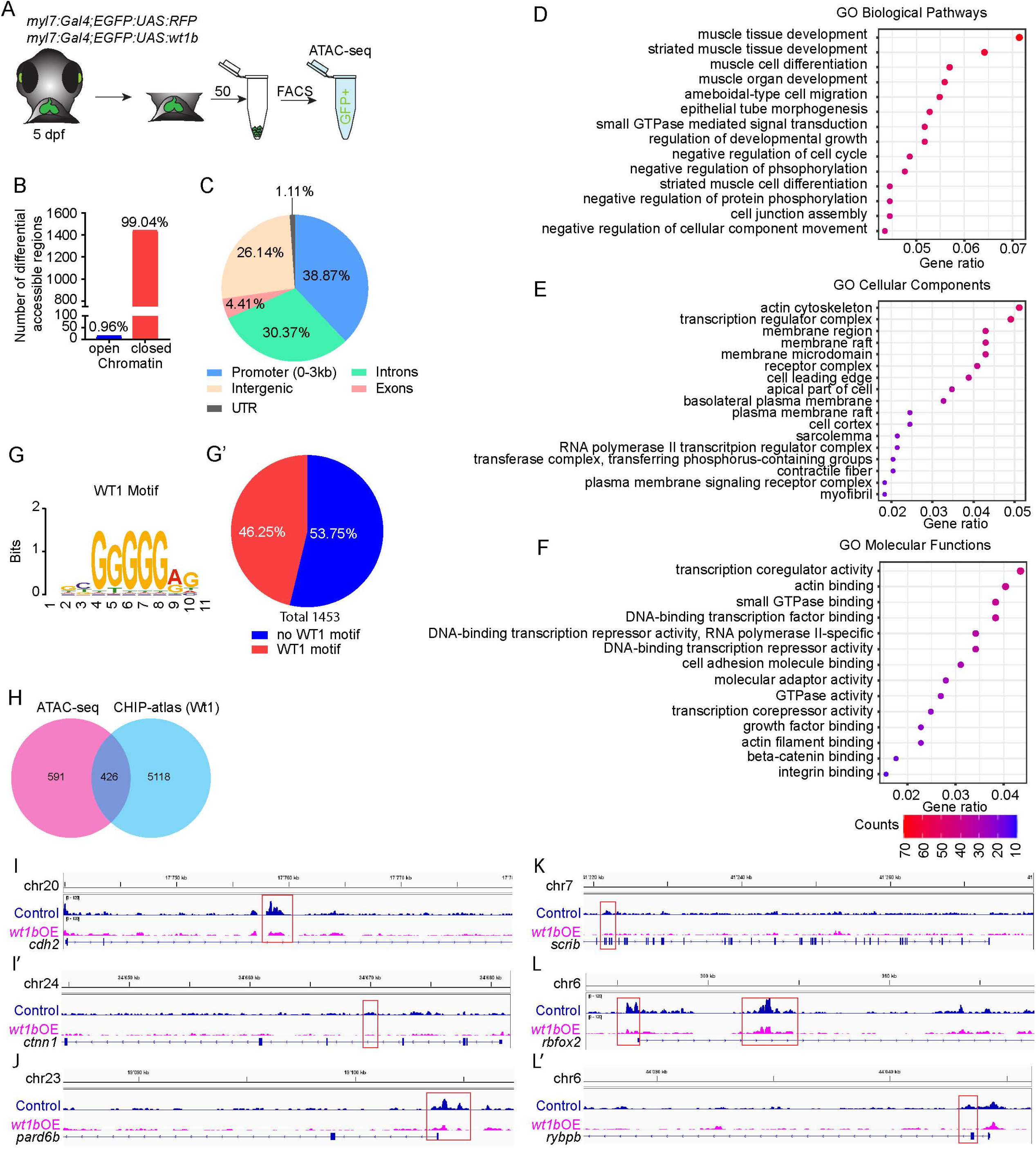
Assay for Transposase-Accessible Chromatin sequencing (ATAC-seq) in *wt1b*-overexpressing cardiomyocytes. (A) Schematic representation of cell acquisition for ATAC-seq. (B) Graphical representation of number of differential accessible regions between *myl7:Gal4; eGFP:UAS:RFP* and *myl7:Gal4; eGFP:UAS:wt1b* cardiomyocytes at 5 days post-fertilization (dpf). (C) Distribution of the genomic regions with differential accessible regions. (D-F) Gene Ontology (GO) pathways enrichment for differential accessible regions in cardiomyocytes after *wt1b*-overexpression. (D) Shown are selected GO Biological Pathways enrichment out of the top 25. (E) Shown are selected GO Cellular Components enrichment out of the top 25. (F) Shown are selected GO Molecular Functions enrichment out of the top 25. The color scale indicates the number of genes enriched in a pathway. All pathways have enrichments significance p-adjust ≤0.05. (G-G’) MEME-Centrimo WT1 motif analysis. (G’) Percentage of the differential accessible regions in which the WT1 motif is represented. (H) Venn Diagram comparing the number of differential accessible regions that are common between the ATAC-seq and the CHIP-atlas database for WT1. (I-L’) Sequencing tracks for genes with differential peaks within their genomic loci. Shown are genes representative of adherens junctions: *cdh2* (I) and *ctnn1* (I’); apical polarity, *pard6b* (J); basal polarity, *scrib* (K); and sarcomere assembly: *rbfox2* (L) and *rybpb* (L’).

Having seen that overexpression of *wt1b* in cardiomyocytes affected heart development and that these changes correlated with the observed molecular signature, we looked more closely on how the genetic landscape of some of the genes with associated differentially accessible regions was affected. We had previously seen that apical cell-cell junctions were disrupted in the hearts of the *myl7:Gal4;eGFP:UAS:wt1b* embryos, including expression and localization of *cdh2*, ZO-1, β-Catenin. In agreement, we observed that putative regulatory regions near *cdh2* and *ctnna* (another core component of adherens junctions) revealed lower accessibility when *wt1b* was overexpressed in cardiomyocytes (Fig 6I-I’). We also observed lower accessibility in core apicobasal polarity pathway genes (47) such as *pardb6* and *pard3bb* from the apical polarity pathways and *scrib*, *dlg1* and *dlg1l* from the basolateral pathways (Fig 6J-K, S3 Data), supporting a perturbed apicobasal polarity in the *wt1b* overexpression lines. Moreover, we detected that several genes associated with sarcomere assembly such as e2f3, rbfox2 and rybp (48–50) presented lower chromatin accessibility in *wt1b*-overxpressing cells (Fig 6L-L’, S3 Data), which could explain the disrupted sarcomeres observed in the overexpression line (Fig 5M-R’).

In conclusion, ATAC-seq data analysis revealed that *wt1b* overexpression in the heart decreased overall chromatin accessibility associated with key genes involved in cardiomyocyte maturation and structural differentiation, with Wt1b likely to directly repress gene expression programs controlling muscle development, cell polarity and actin binding.

### Overexpression of *wt1* in cardiomyocytes during embryogenesis impairs heart morphogenesis and induces fibrosis in the adult heart

In *wt1b*-overexpression hearts, several of the top enriched Biological Pathways were associated with muscle development. Moreover, these hearts showed several impairments in cardiomyocyte differentiation and fate. In view of this, we decided to take a closer look at the overall changes in cardiac morphology and growth upon sustained myocardial *wt1b/a* overexpression. We had previously noted that those animals with strong eGFP expression throughout the myocardium presented impaired cardiac looping (Fig 5F-F’), often with a heartstring morphology (observed in n=5 out of 5 embryos by whole mount immunofluorescence). We performed *in vivo* imaging between 2 and 3 dpf, the time window of cardiac looping (4). We found that, whereas the heart of a *myl7:Gal4;eGFP:UAS:RFP* embryo looped normally, in a *myl7:Gal4;eGFP:UAS:wt1b* larva the heart started to loop, but eventually this process stopped and reverted, resulting in a tubular-like shaped heart (S6 Fig and S8 Video; n=2). We analyzed looping dynamics by quantifying the angle between the ventricle and the atrium (51, 52) (S6 Fig). Whereas in 5 dpf control hearts the angle between the ventricle and the atrium was, on average, lower than 110° (108°±5), in *wt1b*-overexpressing hearts the angle was larger (142°±15) (S6 Fig).

To validate that these morphological changes were specific to the overexpression of *wt1* in cardiomyocytes we decided to induce *wt1b* expression in other cardiac cell populations (S3 Fig). For that, we crossed the *Tg*(*eGFP:UAS:wt1b)* into *Tg*(*fli1a:Gal4)* (51), to overexpress *wt1b* in the endocardium (S3 Fig), and into *TgBAC(nfatc1:GAL4ff)^mu286^* (52), to overexpress *wt1b* in the atrioventricular valves (S3 Fig). We could not detect any apical delamination, looping defects or reduced MHC expression in these hearts at 3 dpf and 5 dpf.

As cardiomyocyte and general heart morphology were affected, we decided to evaluate cardiac performance. We did *in vivo* imaging and analyzed different parameters for heart function in *myl7:Gal4;eGFP:UAS:wt1b* and *myl7:Gal4; eGFP:UAS:RFP* larvae. We analyzed cardiac function at 2 dpf, the time point at which we first observed cardiac malformations, as well as at 5 dpf, once looping has concluded (S6 Fig; n=14). First, we assessed stroke volume (53, 54), which indicates the volume of blood that the heart is capable of pumping in each contraction. *myl7:Gal4;eGFP:UAS:wt1b* ventricles presented a reduced stroke volume at 2 dpf (0.11±0.04 nl *vs* 0.04±0.03 nl) and this impairment did not recover at 5 dpf (0.39±0.17 nl *vs* 0.22±0.08 nl) (S6 Fig). We next analyzed the heart rate. Although at 2 dpf we could not detect changes in heart rate (114±8 beats per min (bpm) *vs* 119±8 bpm) we observed a significant decrease in the *wt1b* overexpression heart frequency at 5 dpf (166±13 bpm *vs* 141±9 bpm) (S6 Fig). The reduced stroke volume together with the decreased heart rate indicates that the *wt1b* overexpression animals have also an impaired cardiac output. Following on this observation, next we measured the ejection fraction for the ventricle and the atrium (54). We found that in the atrium, at 2 dpf, the ejection fraction did not significantly change between both groups (43±14 % *vs* 51±8 %). However, at 5 dpf there was a clear reduction in the ejection fraction of the atrium (55±8% *vs* 41±12%). In contrast, the ventricular ejection fraction was initially significantly reduced at 2 dpf in *wt1b* overexpressing embryos (48±13% vs 35±13%), but recovered at 5 dpf (50±8% vs 49±8%) (S6 Fig).

We also noted that, already at 2 dpf, the atria of *wt1b-*overexpressing animals seemed to be much larger than that of the *eGFP:UAS:RFP* line (Fig 7A-C’). This difference in atria size was sustained at 5 dpf (Fig 7D-E’). To confirm that the atria were indeed larger in these animals we first calculated the ratio between the atrium and ventricle volume at 5 dpf. We saw that in *wt1b*-overexpressing hearts at 5 dpf the atria were on average 1.5 times larger than the ventricle, whereas in the control group they were only 0.5 times bigger (Fig 7F). To evaluate what could be the cause of this atrial enlargement, we counted the number of atrial and ventricular cardiomyocytes. While the number of MHC-positive cells in the ventricles was only slightly smaller than in the *wt1b*-overexpressing hearts (163±47 *vs* 95±24), there was a significant increase in MHC-positive cells in *myl7:Gal4;eGFP:UAS:wt1b* atria (63±8 *vs* 132±11) (Fig 7 G-G’). This indicated that atrial enlargement in *wt1b*-overexpressing hearts might be due to cell hyperplasia. To understand what could be the source of the excess of atrial cells, we performed BrdU staining to evaluate proliferation. We calculated the ratio of proliferating cardiomyocytes per total amount of cardiomyocytes for each chamber. Contrary to our expectations, in the atria of *wt1b*-overexpressing hearts cardiomyocyte proliferation was significantly reduced (Fig 7H), while in the ventricle proliferation was not affected (Fig 7H’). This could indicate that atrial hyperplasia is more likely to be due to a continuous inflow of cardiac precursors, rather than overproliferation of the cells in this chamber. Atrial enlargement persisted also in juvenile stages (S7 Fig). Due to the severity of the phenotype (S2 Table), few *myl7:Gal4;eGFP-UAS-wt1b* animals survived past 6 dpf. Therefore, we used the *myl7:CreERT2:eGFP-T2A-wt1a* line to evaluate the morphology of the adult heart. We analyzed adult hearts from embryonically recombined and 4-OHT untreated *myl7:CreERT2:eGFP-T2A-wt1a* animals (Fig 7K). Consistent with our result with *wt1b*, we observed that animals overexpressing *wt1a* in cardiomyocytes starting at an embryonic stage, revealed an enlarged atrium (22/42) (Fig 7 L-M). The increase could be quantified by micro computed tomography scanning (micro-CT) and shown to correspond to a doubling of the normal atrial volume (Fig 7N-R”).

**Fig 7.**
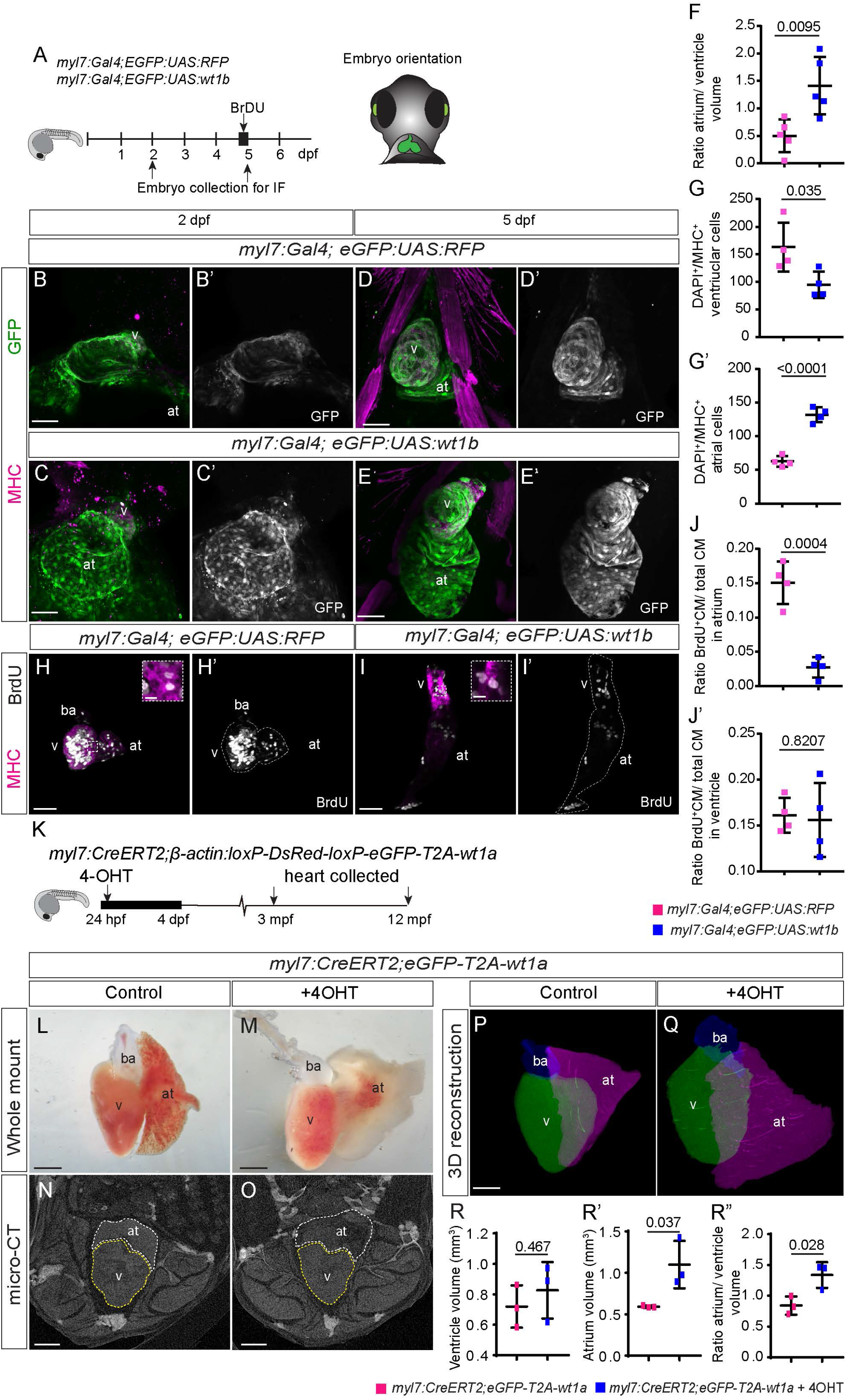
Overexpression of *wt1a* or *wt1b* in cardiomyocytes causes morphological changes in the zebrafish heart. (A) Schematic representation of lines used and embryo orientation for imaging. (B-E’) Immunofluorescence against GFP and myosin heavy chain (MHC) on *myl7:Gal4; eGFP:UAS:RFP* and *myl7:Gal4;eGFP:UAS:wt1b* zebrafish embryos. Shown are 3D projections of the heart region of 2 days postfertilization (dpf) embryos (B-C’), 6 dpf larvae (D-E’). (F) Quantification of the ratio of the atrium and ventricle volumes of 5 dpf zebrafish hearts. Statistical significance was calculated by unpaired t-test. Shown are mean ±SD as well as individual measurements. (G-G’) Quantification of the number of ventricular (G) and atrial (G’) cardiomyocytes in 5 dpf zebrafish hearts. Statistical significance was calculated by unpaired t-test. Shown are mean ±SD as well as individual measurements. (H-I’) Immunofluorescence against BrdU and myosin heavy chain (MHC) on *myl7:Gal4; eGFP:UAS:RFP* (H-H’) and *myl7:Gal4;eGFP:UAS:wt1b* (I-I’) zebrafish embryos. Shown are 3D projections of the heart region of 5 dpf. The box on the top right corner of panels H and I show zoomed views of the boxed region in the 3D projections of the hearts. Zoomed views are maximum intensity projections of two consecutive slices highlighting the BrdU and MHC positive cells. (J-J’) Quantification of the ratio of the BrdU+ cardiomyocytes per total number of cardiomyocytes in the atrium (H) and ventricle (H’) of 5 dpf zebrafish hearts. Statistical significance was calculated by unpaired t-test. Shown are mean ±SD as well as individual measurements. (K) Schematic representation of the time points during which 4-hydroxy-tamoxifen (4-OHT) was administered to *myl7:CreERT2;β-actin:loxP-DsRed-loxP-eGFP-T2A-wt1a* fish (in short *myl7:CreERT2;eGFP-T2A-wt1a)*. Controls are *myl7:CreERT2;eGFP-T2A-wt1a* that were not treated with 4-OHT. Hearts were collected at 12 months postfertilization (mpf). (L-M) Bright field images of whole mount adult zebrafish hearts untreated (L) and treated with 4-OHT during embryogenesis (M). (N-O) micro-computed tomography (micro-CT) image of adult heart of untreated (N) and fish treated with 4-OHT (O) during embryogenesis. (P-Q) 3D volumetric rendering of 3D images acquired with a microCT of adult hearts. (P) untreated, (Q) 4-OHT treated. (R-R’) Quantification of chambers volumes of adults *myl7:CreERT2;eGFP-T2A-wt1a* that were not treated with *4-OHT* hearts. (R) Shown are the differences in ventricle volume between recombined and non-recombined hearts. Each point represents one heart. (R’) Quantification of the differences in atrium volume between recombined and non-recombined hearts. Each point represents one heart. (R”) Quantification of the ratio between the volume of the atrium and the ventricle from micro-CT images acquired from heart of the two experimental groups. Each point represents one heart. Statistical significance was calculated with an unpaired t-test. Shown are means ±SD. Scale bar: 50 µm (B-E’ and H-I’) and 500 µm (L-Q). at, atrium; v, ventricle; ba, bulbus arteriosus.

We further analyzed *myl7:CreERT2; eGFP-T2A-wt1a* hearts on histological sections (S7 Fig). Similarly to what we had seen in the juvenile hearts of the *wt1b* overexpressing fish (S7 Fig), we found a high degree of myocardialization of *wt1a*-overexpressing atria, a feature resembling trabeculation in the ventricle (S7 Fig, n=3/4). Furthermore, we detected the deposition of fibrotic tissue around atrial walls (S7 Fig, n=4/4). Immunolabelling with anti-Col1a1 confirmed these findings. Whereas in the control animals Col1a1 labelling was only detected in the valves (S7 Fig), in hearts of recombined *myl7:CreERT2; eGFP-T2A-wt1a* animals large regions of the atria were also Col1a1-positive. These were in close proximity with eGFP-positive cells, which might indicate that *wt1a*-expressing cardiomyocytes are secreting Col1a1 (S7 Fig). In sum, induced expression of *wt1a/b* in cardiomyocytes leads to atrial hypertrophy, which in the adult is accompanied by interstitial fibrosis.

Taken together, our data indicates that apart from the induction of cell fate change from cardiomyocytes to epicardial cells, overall, sustained expression of *wt1b* in cardiomyocytes affects heart development leading to impaired cardiomyocyte maturation, increased atrial size due to cardiomyocyte hyperplasia, as well as defective cardiac looping and heart function.

## DISCUSSION

During myocardial development, cells from the precardiac mesoderm enter the heart tube. In the zebrafish, the myocardial tube is comprised of an epithelial lining which forms a continuum with the *wt1a* and *wt1b*-positive pericardial mesothelium (9, 18). We observed that during heart tube extension, *wt1b*-positive mesothelial cells enter the heart tube and differentiate into cardiomyocytes. Concomitantly, *wt1b* reporter gene expression is downregulated, suggesting that wt1 downregulation is needed for myocardial maturation. This process seems to be conserved across species as active repression of wt1 locus is detected also detected during the differentiation process of mouse embryonic stem cells into cardiomyocytes (20). In addition, recently, it has been shown that during early stages of mouse heart development there is a common progenitor pool that can give rise to both epicardial as well as myocardial cells (55). Given that sustained wt1 activity reduced chromatin accessibility in regulatory regions associated with cardiomyocyte–specific genes and that wt1 activity in cardiomyocytes can induces their phenotypic switch from myocardial to epicardial cells, we conclude that wt1 downregulation is a prerequisite for cardiomyocyte differentiation.

During proepicardium formation, *wt1*-positive cells apically extrude from the dorsal pericardial mesothelium giving rise to proepicardial cell clusters that subsequently are transferred to the myocardium (56). Here we find that *wt1*-positive cells in the myocardium undergo a similar process and delaminate apically from the myocardial epithelium. It will be important to further decipher possible parallelisms between these two processes and elucidate the direct role of *wt1* during these cellular rearrangements. Wt1 participates in the mesothelial-to-mesenchymal transition giving rise to epicardial derived cells (EPDCs) (57, 58). Moreover, Wt1 has been suggested to control the retinoic acid (RA) signaling pathway during EPDC formation (58),(59). The fact that cardiomyocytes overexpressing *wt1* are relocating to the epicardial layer might indicate that these cells undergo EMT-like processes in response to *wt1* overexpression, a process, which might be mediated by RA. However, we did not observe *aldh1a2* expression in the myocardium, prior to delamination suggesting that *aldh1a2* expression might be a consequence rather than a cause of apical delamination of *wt1a* or *wt1b* expressing cells. Of note, not all eGFP-positive cardiomyocytes undergo delamination. It might thus be possible that not all cardiomyocytes have the capacity to respond to the same extent to *wt1* overexpression. Indeed, in the mouse a small subset of cardiomyocytes has been shown to express *Wt1* and as such, not all cardiomyocytes might be equally sensitive to a change in Wt1 dosage (60, 61).

Cardiomyocyte extrusion has been observed in *klf2* and *snai1* mutant zebrafish (26, 62)). While in both cases, extruded cardiomyocytes are eliminated, here we report that the extruded cells remain on the myocardial surface contributing to the epicardial layer.

*Wt1* lineage tracing studies using Cre/lox transgenic lines in the mouse, suggested that epicardial derived cells were able to contribute to cardiomyocytes during development and repair (63, 64). Here we report the opposite phenotypic switch induced by Wt1. *Wt1* overexpression in cardiomyocytes had also been suggested to trigger a change in cell fate in a pathological condition (65). In arrythmogenic right ventricular cardiomyopathy (ARVC), a disease-causing arrhythmia leading to the accumulation of fat deposits in the heart, a subset of cardiomyocytes has been suggested to start to express *Wt1* and convert into adipocytes. Interestingly, epicardial fat represents an epicardial derivative (66). Together with our results, this indicates that expression of *Wt1* in cardiomyocytes contributes to a phenotypic change, transforming them into epicardial cells or EPDC-like cells. In line with this, in the adult heart we observed the deposition of fibrotic tissue in close proximity to *wt1a*-overexpressing cells. The fibrosis might be a consequence of atrial hypertrophy, that is often accompanied by scar deposition(67), or, alternatively, might indicate that *wt1a* overexpressing cells differentiate or adopt features of EPDCs, in this case extracellular matrix producing fibroblasts.

*wt1b* overexpressing hearts revealed defects such as alterations in muscle cell maturation and sarcomere organization. The fact that sarcomere assembly and stabilization was affected could indicate a general reduced maturity of cardiomyocytes upon overexpression of *wt1b*, which also comes in line with the increased expression of Alcam (43), and reduction in chromatin accessibility of regulatory regions in the vicinity of genes associated with GO pathways related with muscle cell and tissue development and differentiation. Previous work hinted that Wt1 expression prevented the activation of a muscle differentiation program in metanephric-mesenchymal stem cells (68). Also, recent work on the overexpression of *wt1* in an in vitro model of cardiomyocyte differentiation showed reduced cardiomyocyte contractility (69), supporting our observations that wt1 downregulation is a prerequisite to allow myocardial maturation.

A striking phenotypic consequence of *wt1* overexpression is atrial hyperplasia. Enlarged atria might be caused by over proliferation of cardiomyocytes in the atrium, or by increased incorporation of cardiac progenitors from the pericardial mesothelium. Recently it was shown that *lamb1a* mutants have atrial enlargement, most likely due to an excess of second heart field progenitors being added to that region (41). Since we did not observe increased cell proliferation in the atria, the main reason for observing larger atria upon *wt1* overexpression might be that more precursor cells enter the heart during embryogenesis, which might also be linked to the reduced expression of Laminin we observed. This might be secondary to the delay in maturation, which increases the extent of precursors entering the heart.

In conclusion, induced expression of *wt1* in cardiomyocytes during embryogenesis impairs cardiomyocyte maturation and promotes a fate change from cardiomyocytes to epicardial cells. This suggests that during cardiac development, *wt1a/b* expression is turned off in cardiomyocytes once they enter the heart tube to allow their correct differentiation. Dissecting the regulatory mechanisms controlling *wt1a/b* transcription in cardiomyocyte precursors will further expand our knowledge on the tight spatio-temporal control of heart tube expansion and concomitant differentiation.

## Materials and Methods

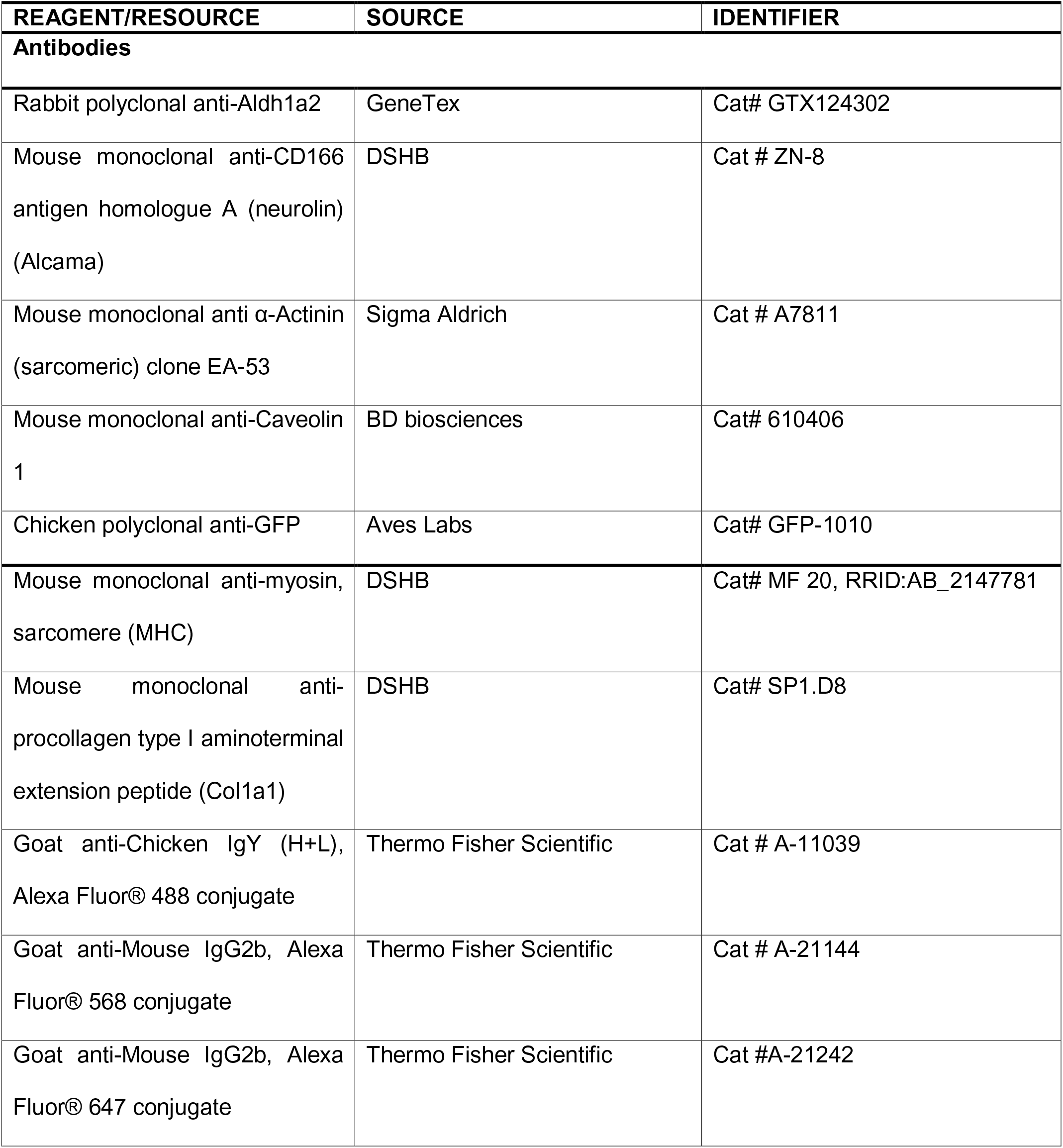
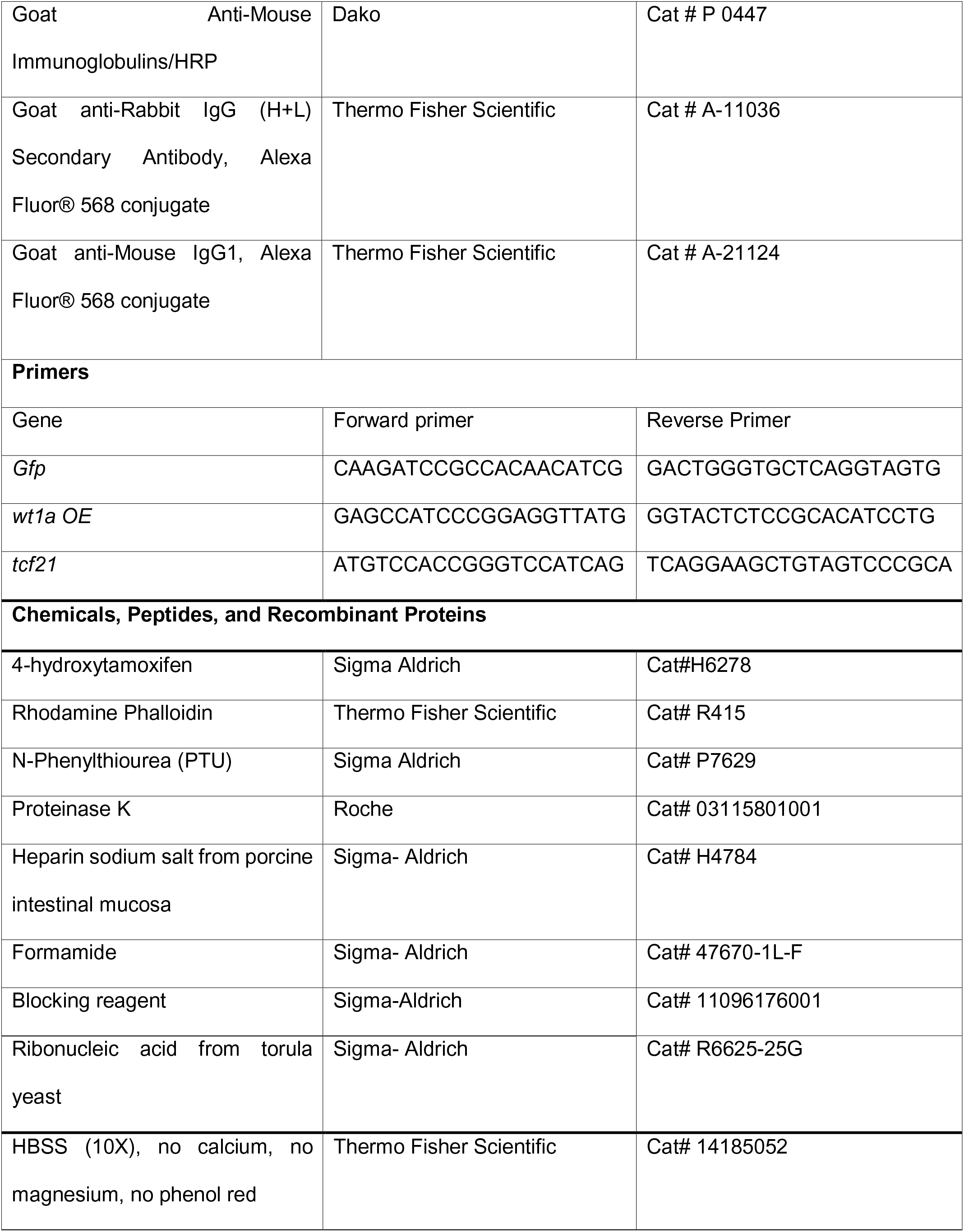
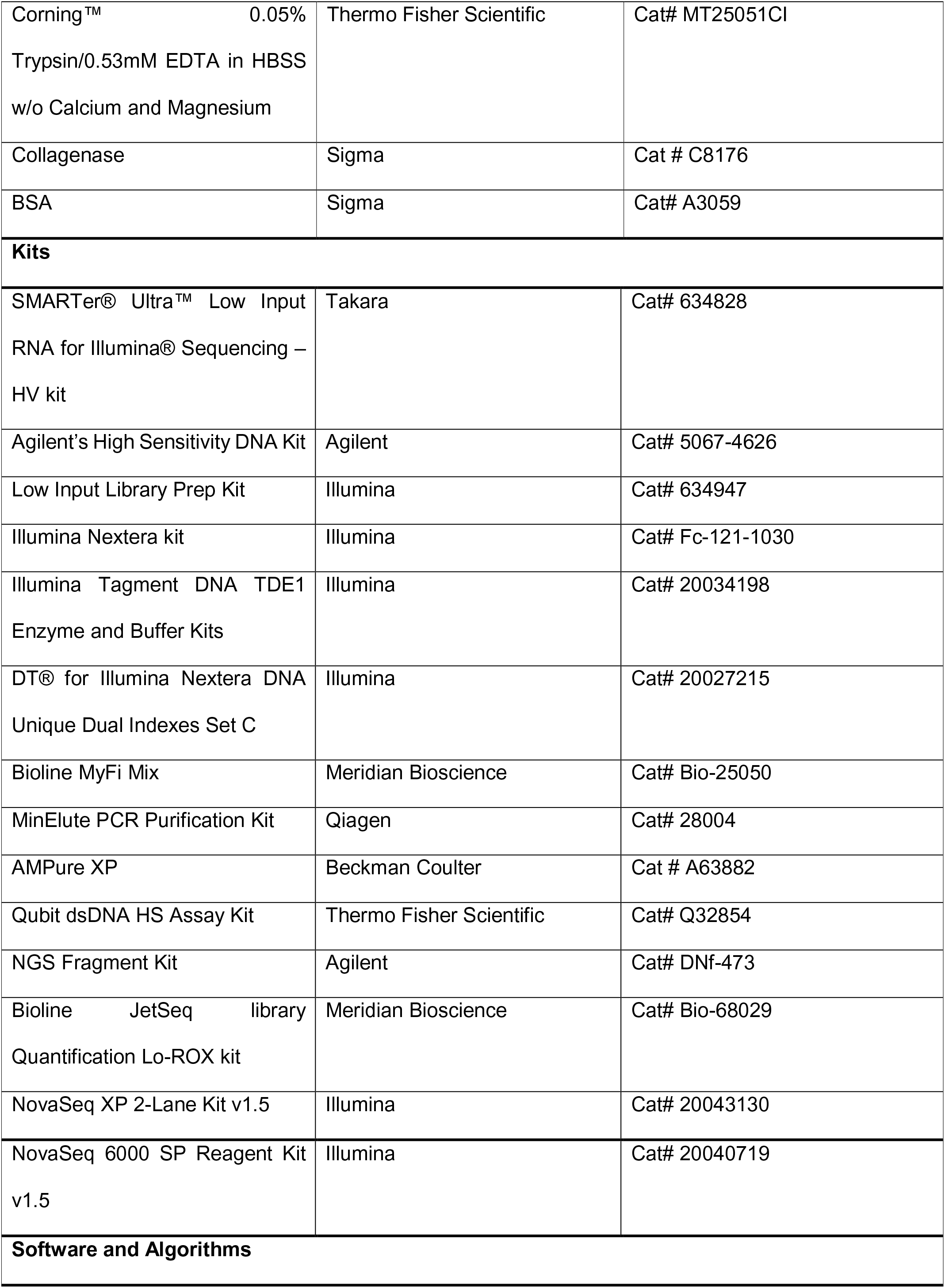
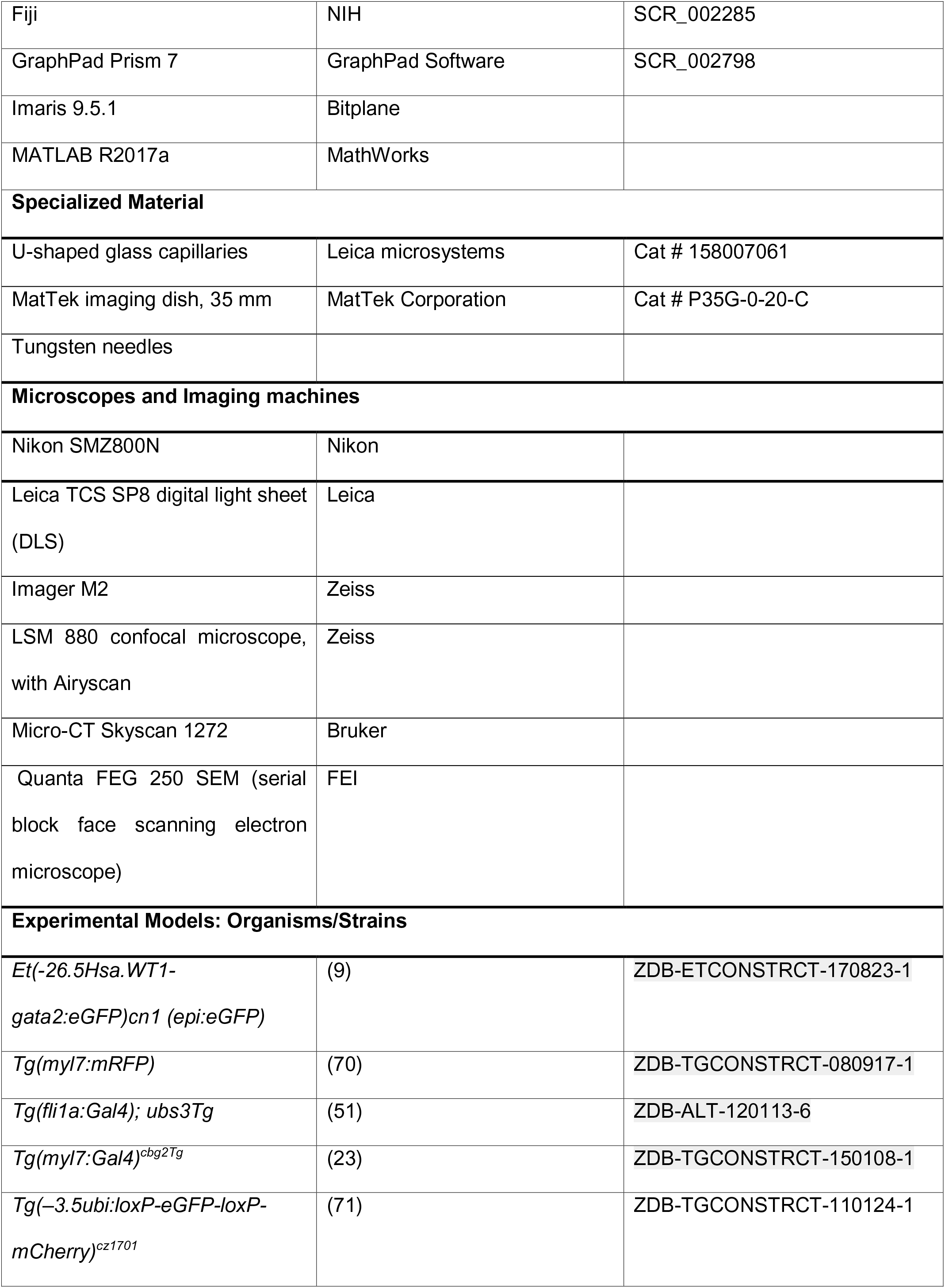
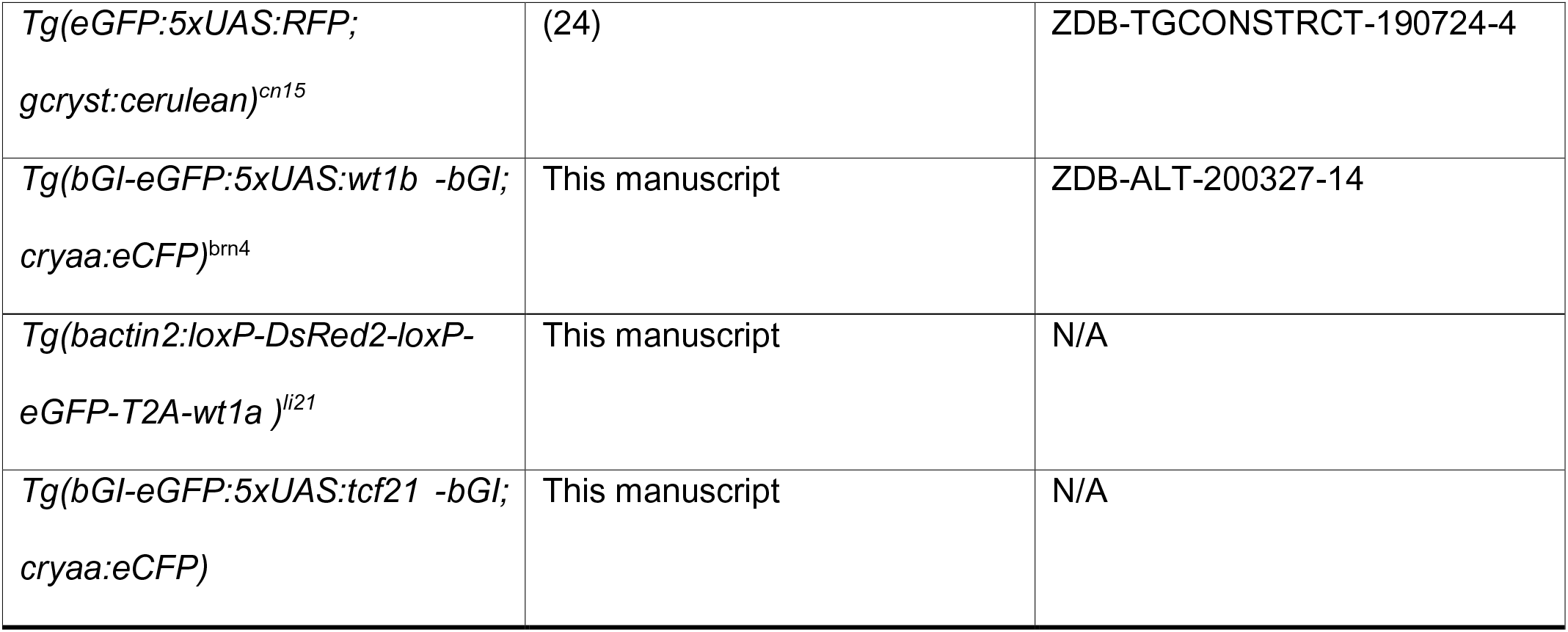

### Zebrafish husbandry

Experiments were conducted with zebrafish embryos and adults aged 3–18 months, raised at maximal 5 fish/l. Fish were maintained under the same environmental conditions: 27.5-28°C, with 14 hours of light and 10 hours of dark, 650-700μs/cm, pH 7.5 and 10% of water exchange daily. Experiments were conducted after the approval of the “Amt für Landwirtschaft und Natur” from the Canton of Bern, Switzerland, under the licenses BE95/15 and BE 64/18.

### Generation of transgenic lines

To generate the transgenic line *eGFP:UAS:wt1b* and the *eGFP:UAS:tcf21* the RFP fragment from the plasmid used to clone *eGFP:5xUAS:RFP* (24) was replaced by either the coding sequence of *wt1b(-KTS)* isoform or of *tcf21*, PCR amplified from 24 hpf and 5 dpf zebrafish embryo cDNA and assembled using Gibson cloning. The final entire construct is flanked with *Tol2* sites to facilitate transgenesis. In this line, tissue specific expression of Gal4 drives the bidirectional transactivation of the UAS leading to the expression of both *eGFP* and *wt1b(- KTS)* or *tcf21* coding sequence. The full name of these lines is *Tg(bGI-eGFP:5xUAS:wt1b(- KTS)-bGI; cryaa:eCFP)*^brn4,^ *Tg(bGI-eGFP:5xUAS:tcf21)-bGI; cryaa:eCFP)*^brn4^.

The construct *bactin2:loxP-DsRed2-loxP-eGFP-T2A-wt1a* was generated by Gateway cloning (MultiSite Gateway Three-Fragment Vector Construction Kit; Invitrogen). As destination vector pDestTol2pA2 was used. The floxed *DsRed2* cassette was derived from vector *pTol2-EF1alpha-DsRed(floxed)-eGFP* (72) and the *wt1a* cDNA was amplified from vector *pCS2P-wt1a* (14). The final construct is flanked with *Tol2* sites to facilitate transgenesis. In the resulting zebrafish line *DsRed* is expressed from the ubiquitous *β-actin* promoter. After Cre-mediated excision of the STOP cassette both *eGFP* as well as *wt1a* are expressed in a tissue-specific manner. The full name of this line is *Tg(bactin2:loxP-DsRed2-loxP-eGFP-T2A-wt1a)^li21^*.

### Administration of 4-Hydroxytamoxifen (4-OHT) to embryos and juvenile fish

4-hydroxytamoxifen (4-OHT; Sigma H7904) stock was prepared by dissolving the powder in ethanol, to 10 mM concentration. To aid with the dissolution the stock was heated for 10 minutes (min) at 65°C and then stored at -20°C, protected from the light. 4-OHT was administered at the indicated times, at a final concentration of 10 µM. For embryos, treatments were performed continuously. For juvenile fish 4-OHT was administered overnight in E3. Prior to administration, the 10 mM stock was warmed for 10 min at 65 °C (73).

### *In vivo* light sheet fluorescence microscopy and retrospective gating

For *in vivo* imaging of the beating zebrafish heart, 2 dpf old embryos were pipetted with melted 1% low melting agarose in E3 medium (about 45°C), containing 0.003% 1-phenyl-2-thiourea (PTU) (*Sigma-Aldrich*) to avoid pigmentation and Tricaine at 0.08 mg/ml, pH 7 to anaesthetize the fish, into a U-shaped glass capillaries (Leica microsystems). This U-shaped capillary was mounted in a 35 mm MatTek imaging dish. The dish was filled with E3 medium containing 0.003% PTU and Tricaine at 0.08 mg/ml, pH 7.

Imaging was performed with the Leica TCS SP8 digital light sheet (DLS) microscope. We used a 25x detection objective with NA 0.95 water immersion and a 2.5x illumination objective with a light sheet thickness of 9.4 µm and length of 1197 µm. The total field of view is 295 x 295 µm, fitting the size of the embryonic zebrafish heart, allowing space for sample drift. The images were acquired in XYTZL-acquisition (XY: single optical section, T: time series, Z: serial optical sections, L: looped acquisition) mode for later retrospective gating. The parameters as shown in Table 3 were applied.

The images were saved as single *.lif*-file and transferred to a workstation (*HP-Z series, Dual Intel Xeon e5-2667 v4 3.2 GHz, 256 GB, NVIDIA GeForce GTX 1080 Ti*). A quality check of the data was performed, before the data were further processed. The survival of the larva until the end of the acquisition, the sample drift and the degree of bleaching were assessed in the *Processor_6D* (https://github.com/Alernst/6D_DLS_Beating_Heart). The data were only used if the larva survived the acquisition. The single *.lif*-file was converted to XYTC *.tif*-files, using the *Converter_6D (*https://github.com/Alernst/6D_DLS_Beating_Heart). Each XYTC file was named in the following format *“Image_R0000_Z0000”* to be recognized for further processing. Retrospective gating was performed as previously described (74–76). The *MATLAB* (R2017a) tool *BeatSync V2.1* was used for retrospective gating (access to the software can be requested from the research group of Michael Liebling). The settings for re-synchronization in the BeatSync software were *“Normalized mutual information”*, *“Recursive Z-alignment”* and *“Nearest-neighbor interpolation”.* One entire heart cycle was re-synchronized in 3D. After re-synchronization, a 3D time lapse of a virtually still heart was created, using the Fiji (77) tool *Make_timelapse* (https://github.com/Alernst/Make_timelapse) using the *Make_timelapse Fiji* plugin. The time lapse was represented as maximum intensity projection or individual optical slices.

### SMARTer-seq

Dorsal pericardium, proepicardium and heart-tube were manually dissected, with tungsten needles, from 60 hpf *epi:eGFP;myl7:mRFP* zebrafish larvae. A minimum of 10 of each tissue/organ were collected for each sample in ice cold PBS. Cells were centrifuged for 7 minutes at 250 g. The excess liquid was removed, and the cells were stored at -80°C until further use. RNA was directly transformed and amplified into cDNA from the lysed tissue using the SMARTer® Ultra™ Low Input RNA for Illumina® Sequencing – HV kit. cDNA quality control was verified using the Agilent 2200 BioAnalyzer and the High Sensitivity DNA Chip from Agilent’s High Sensitivity DNA Kit. Next, 50 ng of amplified cDNA were fragmented with the Covaris E220 (Covaris) and used for preparing sequencing libraries using the TruSeq RNA Sample Prep Kit v2 kit (Illumina), starting from the end repair step. Finally, the size of the libraries was checked using the Agilent 2200 Tape Station DNA 1000 and the concentration was determined using the Qubit® fluorometer (Thermo Fisher Scientific). Libraries were sequenced on a HiSeq 2500 (Illumina) to generate 60 bases single reads. FastQ files were obtained using CASAVA v1.8 software (Illumina). NGS experiments were performed in the Genomics Unit of the CNIC.

### SMARTer-seq analysis

All bioinformatics analysis were performed using bash scripts or R statistical software. Quality check of the samples was performed using FASTQC and reports summarized using MultiQC (78, 79). Adapters from the fastq files were trimmed using fastp software (80). Reads were aligned to GRCz11 danRer11 v102 assembly from Ensembl using STAR (81). The reads were summarized using featureCounts (82). The counts data were imported to Deseq2 and genes who had expression across all samples (rowSums) greater than or equal to 10 were kept ensuring the realiable expression estimates (83). After evaluation of the PCA, one of the samples from the heart tube was determined as an outlier and removed from the downstream analysis. The differential expression analysis was performed using ‘ashr’ LFC Shrikange (84). A gene was considered as significant if the p adjusted value was <0.05. The plots were plotted using ggplot2 (85).

### Immunofluorescence

Whole mount immunofluorescence on embryos was done as previously described (24). Shortly, embryos were fixed over-night, at 4 °C in 4% paraformaldehyde (PFA) (EMS, 15710). Then they were washed with PBS-Tween20 (0.1%) and permeabilized for 30 to 60 min with PBS-TritonX100 (0.5%), depending on the stage and the antibody used. Permeabilization was followed by blocking for 2 hours with histoblock (5% BSA, 5% goat serum, 20mM MgCl2 in PBS). Afterwards, embryos were incubated overnight, at 4 °C, with the primary antibodies, in 5%BSA. The next day embryos were washed with PBS-Tween20 (0.1%) followed by and overnight incubation in the secondary antibodies, at 4 °C, in 5% BSA. Finally, embryos were washed with PBS-Tween20 (0.1%) and a nuclear counterstain with DAPI (Merck, 1246530100) 1:1000 was done.

Immunofluorescence on paraffin sections was performed as previously described (86). Briefly, paraffin sections were dewaxed and rehydrated through a series of ethanol incubations, similar to previously described for histological staining. Afterwards, epitope was recovered by boiling the samples in 10mM citrate buffer, pH 6, for 20 min. Next the same procedure was applied as described above for whole mount immunofluorescence.

The following antibodies were used: primary antibodies - anti-eGFP (Aves, eGFP-1010) was at 1:300, anti-Myosin Heavy Chain at 1:50 (DSHB Iowa Hybridoma Bank, MF20), anti-Aldh1a2 at 1:100 (Gene Tex), anti-Alcam at 1:100 (DSHB Iowa Hybridoma Bank, Zn-8), anti-α-actinin at 1:200 (Sigma Aldrich), anti-embryonic Cardiac-Myosin Heavy Chain at 1:20 (DSHB Iowa Hybridoma Bank, N2.216), anti-Caveolin 1 at 1:100 (BD Biosciences) and anti-Procollagen Type I at 1:20 (DSHB Iowa Hybridoma Bank, SP1.D8). Secondary antibodies were Alexa Fluor 488, 568, 647 (Life Technologies) at 1:250 and biotin anti-rabbit (Jackson Immuno Research, 111-066-003) followed by StreptavidinCy3 or Cy5 conjugate (Molecular Probes, SA1010 and SA1011) at 1:250.

### qRT-PCR assay

Hearts from *Tg(eGFP:5xUAS:wt1bOE-KTS;myl7:Gal4)* and *Tg(eGFP:5xUAS:RFP; myl7:Gal4)* were extracted at 40 dpf. Ventricle, atrium and bulbus arteriosus were manually dissected and stored separately in pools of 5. For each sample, between 3 and 7 biological replicates were collected. Total RNA was extracted by using TRI Reagent (Sigma-Aldrich; Cat-No. T9424) according to the manufacturer’s recommendations. Afterwards, a total of 200 ng of total RNA was reverse-transcribed into cDNA using High Capacity cDNA Archive Kit (Invitrogen Life Technologies; Cat-No. 4374966). Quantitative PCR (qPCR) was performed in a 7900HT Fast real-time PCR system (Applied Biosystems). qPCR was done using Power Up SYBR Green Master Mix (Applied Biosystems, A25742).

PCR program was run as follows: initial denaturation step was done for 30 s at 95°C, followed by 40 cycles at 95°C for 5 s and 60°C for 30 s. To calculate the relative index of gene expression, we employed the 2^-ΔΔCt^ method, using *e1f2α* expression for normalization.

### Double in situ hybridization and immunohistochemistry on paraffin sections

In situ hybridization on paraffin sections was done as follows: paraffin sections were dewaxed and rehydrated through a series of ethanol incubations. Sections were then refixed with 4% PFA at room temperature for 20 min. Afterwards they were washed with PBS and the tissue was permeabilized by incubating the slides with 10 µg/ml of Protease K, for 10 min. at 37°C. Afterwards, slides ware washed with PBS and briefly refixed with 4% PFA. The tissue was then incubated for 10 min with triethanolamine 0.1M, pH8 and 0.25% acetic anhydride. After washing the slides with PBS and RNAse free water the slides were incubated for 3 hours with pre-hybridization buffer (50% formamide, 5X SSC pH 5.5, 0.1X Denhardt’s, 0.1% Tween20, 0.1% CHAPs and 0.05mg/ml tRNA), at 65°C. Afterwards, pre-hybridization buffer was replaced with hybridization buffer (pre-hybridization buffer with mRNA probe). Slides were left to incubate with hybridization buffer overnight at 65°C. The next day, slides ware washed twice with post-hybridization buffer I (50% formamide, 5X SSC pH 5.5 and 1% SDS) and 2 times with post-hybridization buffer II (50% formamide, 2X SSC pH 5.5 and 0.2% SDS). Each wash was done for 30 min at 65°C. Slides were then washed another 3 times with maleic acid buffer (MABT) and then incubated for 1 hour blocking solution (2% fetal bovine serum, heat inactivated, and 1% blocking reagent, in MABT). Tissue was incubated overnight at 4°C with anti-DIG antibody in blocking solution at 1:2000. Finally, sections were thoroughly washed with MABT and incubated in alkaline phosphatase buffer (AP buffer, NaCl 0.1M, MgCl2 0.05M, 10% Tri-HCL pH 9.5). Finally, colorimetric assay was performed using BM purple. After the desired staining was achieved, slides were washed with PBS and fixed with 4% PFA, before mounting them with 50% glycerol and imaged using a Zeiss Imager M2, with and Olympus UC50 camera. After imaging sections were washed and further permeabilized with PBS with 0.5% TritonX-100. Then they were incubated for 2hours with 5% BSA at room temperature and incubated with primary antibody, chicken anti-GFP (1:300 in 5% BSA) overnight at 4°C. The next day slides were washed in PBS-0.1% Tween20 and incubated for one hour at room temperature with secondary antibody anti-chicken-HRP. Signal was obtained by incubating slides with DAB solution for 30 seconds at room temperature. The reaction was stopped with water. Slides were then mounted in 50% glycerol and imaged.

### Whole mount in situ hybridization

Whole mount in situ hybridization was performed as described (87), with some minor adaptations. Embryos were selected at 24 hpf and 3 dpf for eGFP expression. After fixation, the embryos were washed with PBS and gradually dehydrated through a methanol series. Embryos were stored in 100% methanol for a minimum of 2 hours, at -20°C. Afterwards, the embryos were rehydrated and permeabilized with proteinase K (10μg/ml in TBST) at 37°C. Incubation times were adjusted according to the stage of the embryos (24 hpf, 10 min and 72 hpf, 20 min). This was followed by a 20 min incubation in 0.1M triethanolamine (pH 8), with 25μl/ml acetic anhydride.

After 4 hours of pre-hybridization, at 68°C, myl7 riboprobe was diluted in pre hybridization solution, at a concentration of 300ng/ml. The embryos were incubated with the riboprobe overnight, at 68°C. The following day, the riboprobe was removed and the embryos were incubated twice for 30 min with post hybridization solution at 68°C. Embryos were then incubated with blocking buffer, freshly prepared, and afterwards with anti-DIG antibody (in blocking solution), at 1:4000, overnight, at 4°C.

The embryos were then washed extensively with Maleic acid buffer (150mM maleic acid pH 7.5, 300 mM NaCl, 0.1% Tween 20). Finally, the embryos were transferred to a 6-well plate and pre-incubated with AP-buffer (0.1M Tris base pH 9.5, 0.1 M NaCl, 1mM MgCl2, 0.1% Tween 20) and then incubated with BM-purple, at room temperature. As soon as color was visible in the heart of either group (overexpression or control), the staining was stopped in both groups by adding TBST and embryos were re-fixed in 4% PFA.

Using a microscope, we could obtain pictures of the hearts of the embryos. For image acquisition, embryos were mounted on 3% methylcellulose for ease of orientation. Embryos were positioned so that the majority of the heart could be observed in a single plane.

Images were acquired with a Nikon SMZ800N stereomicroscope. Illumination conditions and acquisition parameters were maintained for all embryos.

### Serial block face scanning electron microscopy

Zebrafish embryos at 5 dpf were killed with an overdose of tricaine and immediately fixed with 2.5% glutaraldehyde with 0.15M cacodylate buffer and 2mM CaCl2, pH 7.4. Embryos were then processed for serial block face scanning electron microscopy as previously described (88). Briefly we proceed as follows: samples were rinsed 3 times in ice-cold 0.15 M Na-cacodylate for 5 min. They were then incubated in 0.15 M Na-cacodylate solution containing 2% OsO4 and 1.5% potassium ferrocyanide for 45 min, at room temperature, and for 15 min in a water bath, at 50 °C. Samples were rinsed 3 times for 5 min in water. They were then incubated with 0.64 M pyrogallol for 15 min at room temperature, for 5 min in a water bath at 50 °C, and subsequently rinsed with water. The embryos were incubated in 2% OsO4 for 22 min at room temperature and 8 min in a water bath at 50°C. Afterwards they were again rinsed in water (3 times 5 min) and incubated overnight in a solution of 0.15 M gadolinium acetate (LFG Distribution, Lyon, France) and 0.15 M samarium acetate (LFG Distribution) pH 7.0. The next day the embryos were rinsed 3×5 min with water and incubated in 1% Walton’s lead aspartate (89) at 60 °C for 30 min, and rinsed with water (3x 5 min).

After staining, the samples were dehydrated in a graded ethanol series (20%, 50%, 70%, 90%, 100%, 100%) at 4°C, each step lasting 5 min. They were then infiltrated with Durcupan resin mixed with ethanol at ratios of 1:3 (v/v), 1:1, and 3:1, each step lasting 2 h. The embryos were left overnight to infiltrate with Durcupan. The next day, samples were transferred to fresh Durcupan and the resin was polymerized for 3 days at 60 °C. Sample blocks were mounted on aluminum pins (Gatan, Pleasonton, CA, USA) with a conductive epoxy glue (CW2400, Circuitworks, Kennesaw, GA, USA). Care was taken to have osmicated material directly exposed at the block surface in contact with the glue in order to reduce specimen charging under the electron beam. Pyramids with a surface of approximately 500 × 500 μm2 were trimmed with a razor blade.

Three-dimensional (3D) ultrastructural images were produced by serial block face scanning electron microscopy (SBFSEM) on a Quanta FEG 250 SEM (FEI, Eindhoven, The Netherlands) equipped with a 3View2XP in situ ultramicrotome (Gatan). Images were acquired in low vacuum mode (40 mPa), except where indicated otherwise. Acceleration voltage was 5 kV and pixel dwell time was set between 2 µs. Image acquisition was done with a back scattered electron detector optimized for SBFSEM (Gatan). Image stack were aligned, normalized, and denoised by non-linear anisotropic diffusion in IMOD (90). Each field of view consisted of 8192 x 8192 pixels with a dimension of 6 nm/pixel in x-y and zz nm in z direction. Final image montage was done in Fiji.

### FAC sorting

*myl7:Gal;eGFP:UAS:RFP* and *myl7:Gal;eGFP:UAS:wt1b* embryos at 5 dpf were used to obtain GFP+ heart cells. The heart region of these embryos was manually dissected and placed in Ringer’s solution. Afterwards the tissue was briefly centrifuged in a table top centrifuge and the Ringer’s solution was replaced by a mix of 20mg/ml collagenase in 0.05% trypsin. The samples were incubated at 32°C for 25 minutes. Every 5 minutes this mixture as gently mixed. The tissue was visually inspected for dissociation. After cell disaggregation the reaction was stopped with Hanks’s solution (1xHBS, 10mM Hepes and 0.25%BSA). The homogenized samples were centrifuged at 250g for 10 minutes and re-suspended in Hank’s solution. The cells were then passed through a 40 µm filter, centrifuged gain for 10 minutes at 400g and re-suspended in 50 µl of Hank’s solution for FAC sorting. Dead cells were marked with 7-aminoactinomycin D (Invitrogen) and discarded. Cells were FAC sorted into Hank’s solution, on a Moflo astrios EQ (Beckman Coulter) and analyzed for forward and side scatter, as well as eGFP fluorescence. Between 1200 and 1500 cells per sample were sorted for

### ATAC-seq

FAC sorted GFP^+^ cells were gently centrifuged and Hank’s solution was replaced by lysis buffer (10mM tris-HCL, pH 7.4, 10mM NaCl, 3mM MgCl2, 0.1% IGEPAL CA-630). Cells were immediately centrifuged at 500 g for 10minutes at 4°C. The supernatant was discarded and replaced with the transposition reaction mix (Tn5 in TD buffer) for tagmentation, and incubated at 37°C for 30min. Afterwards, 500mM of EDTA was used for quenching. The solution was incubated for 30 minutes at 50°C. MgCl2 was added to a final concentration of 45mM. Samples were stored at 4°C before proceeding with PCR amplification. PCR amplification we used 1.25µl IDT® for Illumina Nextera DNA Unique Dual Indexes Set C, which contains two indexes premixed and 25 µL of Bioline MyFi Mix. This is in the place of the NEB Next HiFi PCR mix in your protocol. We performed the PCR as outlined. For PCR amplification 15 cycles were used due to the reduced amount of material. The amplified library was purified using the Qiagen PCR purification MinElute kit. This was followed by a 1 x volume AMPure XP bead-based clean-up according to manufacturer’s guidelines. The resulting libraries were evaluated for quantity and quality using a Thermo Fisher Scientific Qubit 4.0 fluorometer with the Qubit dsDNA HS Assay Kit and an Advanced Analytical Fragment Analyzer System using a Fragment Analyzer NGS Fragment Kit, respectively.

The ATAC-Seq libraries were further quantified by qPCR using a Bioline JetSeq library Quantification Lo-ROX kit according to their guidelines. The libraries were pooled equimolar and further cleaned using AMPure XP beads as described above. The library pool was then again assessed for quantity and quality using fluorometry and capillary electrophoresis as described above.

The pool was loaded at 150pM using an XP workflow into one lane of a NovaSeq 600 SP with NovaSeq XP 2-Lane Kit v1.5. The libraries were sequenced paired end on an Illumina NovaSeq 6000 sequencer using a NovaSeq 6000 SP Reagent Kit v1.5. An average of 56 million reads/library were obtained. The quality of the sequencing runs was assessed using Illumina Sequencing Analysis Viewer and all base call files were demultiplexed and converted into FASTQ files using Illumina bcl2fastq conversion software v2.20. ATAC-seq experiments were performed in collaboration with the Genomics Unit of the University of Bern.

### ATAC-seq Data Analysis

All bioinformatics analysis were performed using bash scripts or R statistical software. Quality check of the samples was performed using FASTQC and reports summarized using MultiQC (78, 79). Adapters from the fastq files were trimmed using trimmomatic software (91). Reads were aligned using bowtie2 (92, 93) to GRCz11 danRer11 v102 assembly from Ensembl (81) with flags ‘--very-sensitiv’. Paired-end reads were used for downstream analysis. The files were then converted to bam, downsampled to the lowest counts, indexed and the mitochondrial chromosome was removed using samtools (94, 95). Duplicates were removed using Picard tools (96). The samples were processed to select for unique reads using samtools (94, 95). The peaks were identified using Genrich in ATACSeq mode and zebrafish genome size (97). The zebrafish genome size was estimated using faCount script from public utilities from UCSC (98).

To analyze the differential accessible regions, we used DiffBind using background and DeSeq2 normalization and a cutoff threshold of p<0.05. To annotate the peaks, we used ChIPSeeker (99). We used the GRCz11 danRer11 v102 assembly from Ensembl and transcription start site region as +/-1kb for annotation. The annotated genes were then converted to mouse orthologous genes using biomaRt and used for pathway analysis using clusterProfiler (100, 101). K-means clustering was performed using SeqMINER software using linear enrichment clustering approach with 10 clusters (102). The bigwig files to visualize the peaks were made using bamCoverage in deepTools2 (103). Interactive Genome Viewer was used to visualize the peaks (104).

To identify the transcription factor binding cite, we used the sequences from the differential accessible regions in Centrimo from MEME-suite. We used CIS-BP 2.0 Danio rerio Database to identify the potential zebrafish transcription factors (105, 106).

### Embryonic heart function analysis

Heartbeat analysis was performed by assessing the following parameters: degree of rhythmic beating as Root Mean Square of Successive Differences (RMSSD) (107); stroke volume (SV - difference between diastolic an systolic volume); ejection fraction (EF - difference between diastolic an systolic volume relative to the diastolic size); cardiac output (CO - SV multiplied by heart rate); and diastolic volume, and heart rate as described (54).

We recorded 300 frames of the beating heart in the GFP channel in *Tg(myl7:Gal4; eGFP:UAS:wt1b)* and *Tg(myl7:Gal4; eGFP:UAS:RFP)* at 2 dpf and 5 dpf using the fluorescence stereo microscope Nikon SMZ25 (SHR Plan apo 1x objective, 10x zoom, 2880x2048 pixel, 0.44 µm/pixel, 17 frames/s).

For the analysis of heart function, we defined the volume of the heart, which is calculated by measuring the long diameter (DL) and the short diameter (DS).

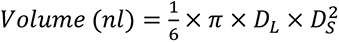

The maximal and minimal volume of the ventricle and atrium were measured, to calculate end-diastolic volume (EDV) and end-systolic volume (ESV). The mean EDV and ESV of two heart cycles per fish were averaged to calculate the SV.

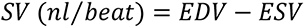

The EF was calculated by dividing the SV through the EDV and converted to a percentage.

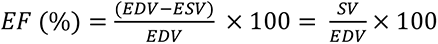

We developed the FIJI plugin *Heart beat analysis* to sequentially process all images in a folder and guide the user through each manual step of the analysis. The manual steps are to find the two diastolic and systolic states of the heart, adjust a line to DL and DS and to draw one line at the border of the ventricle. The plugin *Heart beat analysis* opens subsets of the data (100 frames and only green channel per fish from the *.nd2* RGB file), applies a Gaussian blur filter (10 px), indicates which manual step to perform, calculates the HR by detecting maxima in a kymograph and subsequently saves all kymograph images as *.tiff*, results as *.csv*, all lines as *.zip* in ROI sets.

To calculate the RMSSD we measured the temporal distance between 12-15 cardiac cycles using instead of a subset of 100 frames all 300 frames from the above described data. The temporal distances between cardiac cycles were measured using the FIJI plugin *RMSSD* (107), two line were drawn crossing one side of the cardiac wall of V and AT. Subsequently, kymographs were generated. The correctness of detected maxima in the kymograph was supervised. All intermediate images and ROIs were saved. The locations of each intensity maximum in the kymograph were exported as *.csv*-file. A Jupyter-notebook (108) was created to calculate the time between two cardiac cycles (R – R; time of cardiac cycle, R; current cycle, i; next cycle, i+1) and variability of these time differences between all frames (total frames, N) as RMSSD.

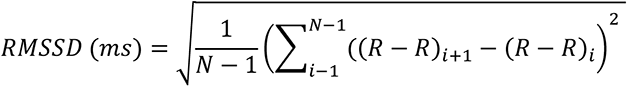

### Histological staining

Hearts were fixed in 2 % paraformaldehyde (PFA) in phosphate-buffered saline (PBS) overnight at 4°C. Samples were then washed in PBS, dehydrated through graded alcohols (30%, 50%, 70%, 90% and 2 x 100%), and Xylol (2x) and embedded in paraffin wax (3x). All steps were done for 20 min. Histological stainings were performed on 8 βm paraffin sections cut on a microtome (Leica and Reichert-Jung), mounted on Superfrost slides (Fisher Scientific), and dried overnight at 37°C. Sections were then dewaxed in Xylol, rehydrated through graded ethanol (from 100% to 30%) and then washed in distilled water. Connective tissue was stained using Acid Fuchsine Orange G (AFOG) (18).

### Micro-computed tomography (microCT)

For each condition, three adult fish were sacrificed in 0.6 mg/ml tricaine (Sigma-Aldrich). Subsequently fish were fixed for 24 hours in 4% paraformaldehyde (PFA), at 4°C. Afterwards, the animals were transversally sectioned bellow the pectoral fins and washed in 1X PBS. They were then stained in lugol for 24 hours, at room temperature, before being scanned by micro-computed tomography. For this, the six samples were imaged on a Bruker SkyScan 1272 high-resolution microtomography machine (Bruker microCT, Kontich, Belgium). The X-ray source was set to a voltage of 80 kV and a current of 125 μA, the x-ray spectrum was shaped by a 1 mm Al filter prior to incidence onto the sample. For each sample, a set of 948 projections of 2452 x 1640 pixels at every 0.2° over a 180° sample rotation was recorded. Every single projection was exposed for 1593 msec. Three projections were averaged to one to reduce the noise. This resulted in a scan time of two hours per sample and an isometric voxel size of 4 μm in the final data sets.

The projection images were then reconstructed into a 3D stack of images with NRecon (Bruker, Version: 1.7.0.4). The 3D images were analyzed using Imaris software and Fiji. For the analysis, the heart and then the atrium, the ventricle and the bulbus arteriosus were segmented and the volume and surface area were obtained. Volume differences between conditions were analyzed using GraphPad Prism7.

For 3D reconstructions of microCT images we used the Fiji software (77, 109). We first selected the images where the heart was visible and created a z-stack with those images. We then proceed to segment three different areas of the heart: the bulbus arteriosus, the atrium and the ventricle. We created a mask in every 5^th^ z-slice and performed a linear interpolation to generate masks for every z-slice. Subsequently, we applied a macro to set all pixels outside of the masked volume to zero. We repeated this process for each one of the three heart areas. We then attributed a different color to each heart region and merged all three channels. This allowed us to represent the segmented parts of the heart and generate a 3D projection (110). We also used the individually segmented heart regions to calculate the volume of each heart chamber. For this we applied the MorphoLibJ plugin (111).

### Imaging and Image processing

Immunofluorescence images were acquired using the Leica TCS SP8 DLS confocal microscopes. For image acquisition of whole mount embryos, larvae were mounted in 1% low melting agarose in a MatTek petri dish. Images were acquired with a 20x water immersion objective. Images were afterwards processed with Fiji software. Fig legends indicate whether a 3D projection is presented or a maximum intensity projection of a reduced number of stacks is shown. For 3D projections, images were first treated with a mean filter, with a radius of 2.0 pixels. Interpolation was also applied when rendering the 3D projections.

To assess the eGFP/mRFP ratio from *in vivo* confocal images, we applied a median filter (3 pixels radius) and measured line profiles from the SV 60 µm into the atrium in 6 sequential Z-planes. The mean intensity along the line profile normalized by the maximum per fluorophore per embryo was calculated, subsequently the ratio for each µm along the line profile was obtained.

Imaging of AFOG stained sections was done with the Zeiss Imager M2, using a 10x objective. For quantification of mean fluorescence intensity first a mean filter with a radius of 2.0 pixels was applied to smoothen the images. Afterwards we did a maximum intensity projection of all the stacks containing the heart. We then delimited the heart and applied an automatic OTSU threshold. Automatic threshold was evaluated independently for each image, when necessary minor adjustments were applied. Finally mean fluorescence intensity was calculated.

Semi-quantification of signal intensity for whole mount in situ hybridization was done using Fiji software. First the images were inverted, region of interest (ROI) was defined and used for all images. For each image mean signal was measured in six independent areas: three in the background and three in the stained area. Measurements were averaged and then background signal subtracted from the signal measured in the stained area. The fold change was calculated and GraphPad was used for statistical analysis.

### Statistical Analysis

Statistical analysis was done with GraphPad Prism 7. When data fitted normality parameters, i.e, passed either the D’Agostino-Pearson or the Shapiro-Wilk normality test, an unpaired t-test was used. If this was not the case, the Mann-Whitney non-parametric test was used to compare differences between conditions. In case a statistically significant difference in the standard deviation between conditions was detected, the Unpaired t test with Welch’s correction was applied. In case of multiple comparisons, a One-Way ANOVA was applied, followed by Tukey’s multicomparisons test. For each graph, in each Fig, the type of statistical test applied is stated in the Fig legend.

The specific test used in each comparison is indicated in the main text or Fig legend. Normal distribution was tested to decide if a parametric or non-parametric test needed to be applied.

### Data Availability

Zebrafish line information has been deposited at ZFIN.

SMARTer-seq and ATAC-seq raw data has been deposited in GEO Database with the reference GSE179520 and GSE179521 respectively.

## ACKNOWLEDGEMENTS

We thank Anna Gliwa and Eduardo Diaz for fish husbandry at University of Bern and CNIC, respectively. Microscopes supported by the Microscopy Imaging Center (MIC) at University of Bern were used. We thank Stephan Müller from the FACSLab of the University of Bern for help with FACS. We thank the Genomics Unit from CNIC for help with SMARTer-seq and Pamela Nicholson and Cátia Coito, from the Genomics Unit of the University of Bern, for help with ATAC-seq. We Thank Didier Stainier for sharing the *Tg(myl7:cdh2-tdTomato)^bns78^* line. NM has been funded by SNF grant 320030E-164245 and ERC Consolidator grant 2018 819717. The CNIC is supported by the Instituto de Salud Carlos III (ISCIII), the Ministerio de Ciencia e Innovación (MCIN) and the Pro CNIC Foundation, and is a Severo Ochoa Center of Excellence (SEV-2015-0505). Benoît Zuber is supported by SNF grant 179520 and ERA-NET NEURON grant 185536. M.O. was supported by SNF grant PCEFP3_186993.

## AUTHORS CONTRIBUTION

I.M. performed most of the experiments, analyzed data, contributed to interpretation of results and wrote the manuscript

A.E. contributed to *in vivo* imaging and image processing and quantifications, contributed to writing the manuscript and interpretation of results

P.A. performed sequencing analysis, contributed to writing the manuscript and interpretation of results

A.V. performed immunofluorescence and helped with embryo dissociation for FACS

T.H. performed qPCR and contributed to other experiments

A.S.-M. generated the *eGFP:UAS:wt1b* line

U.N. generated the *Tg(bactin2:loxP-DsRed2-loxP-eGFP-T2A-wt1a)* line

A.O. performed electron Microscopy imaging and image reconstruction

X.L. contributed to histological staining, sectioning and maintenance of lines

L. A.D. contributed to Smart-Seq

D. H. performed micro-CT imaging and imagine reconstructions

B. Z. supervised electron Microscopy imaging and image reconstruction

R. H. supervised micro-CT imaging and imagine reconstructions

C.T. performed data analysis not included, but with impact to this work

M.O. provided Fig1 S1 and the interpretation thereof.

F.S helped with ATACseq generation and interpretation thereof

C.E. supervised the generation of the *Tg(bactin2:loxP-DsRed2-loxP-eGFP-T2A-wt1a)*

N.M. conceived the research question to be addressed, contributed to design experiments and interpretation of results, wrote the manuscript, and secured funding.

## CAPTATION SUPPORTING FILES

**S1 Fig.**
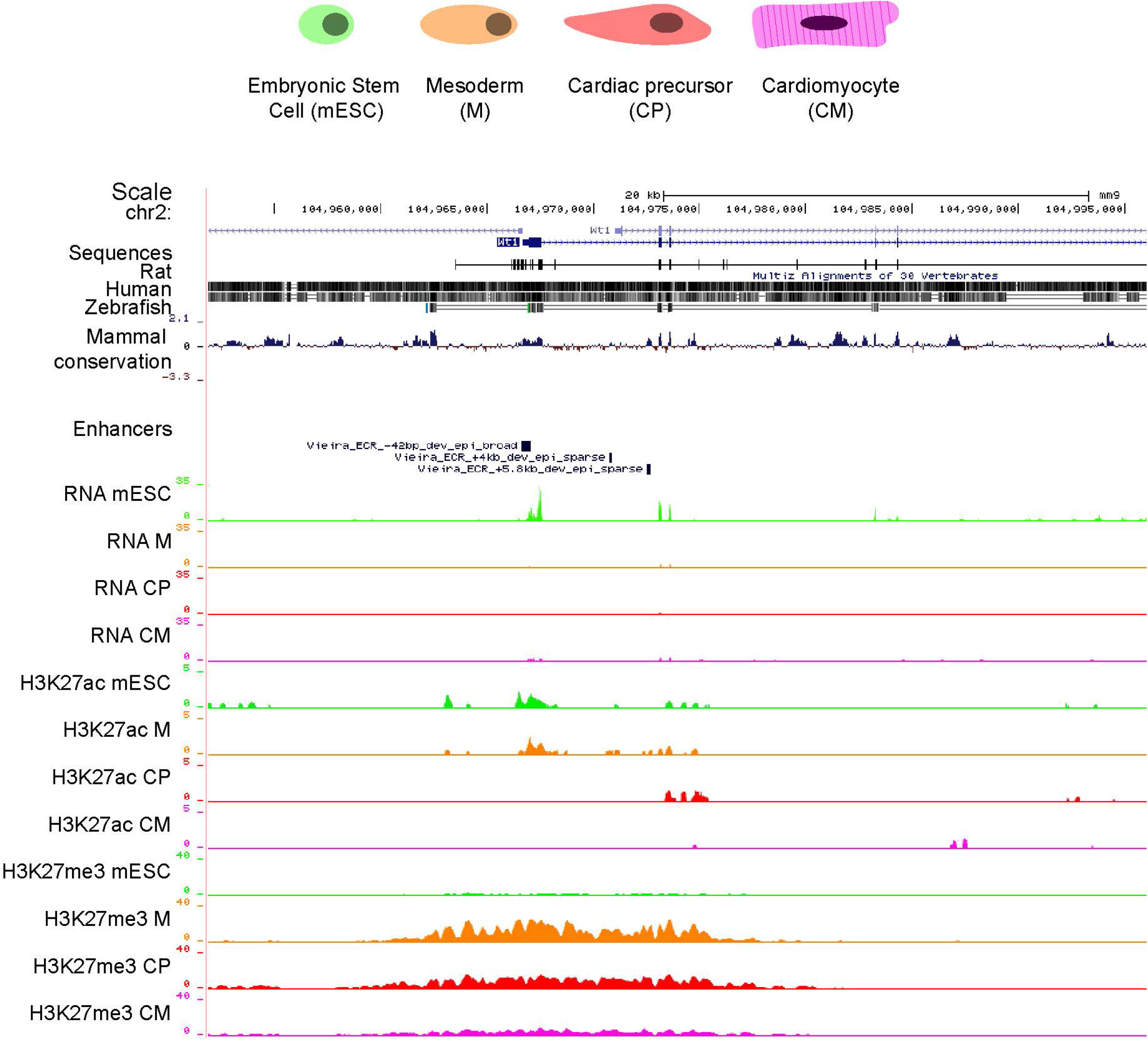
Transcript and histone modification signatures at the *Wt1* locus during *in vitro* differentiation of mouse embryonic stem cells into cardiomyocytes. UCSC browser window depicting the *Wt1* locus and associated transcriptomic and epigenomic signatures in mouse embryonic stem cells (mESC), mesodermal progenitors (M), cardiac precursors (CP) and cardiomyocytes (CM), as published previously (20). Tracks of activating H3K27ac and repressive H3K27me3 marks are shown and co-localizing elements with previously validated epicardial enhancer activities 4kb and 5.8kb downstream of the *Wt1* transcriptional start site (TSS) are indicated (21). Shown tracks represent the sum of the tracks for the different samples, for each type of cell. Mammal conservation is illustrated by the Placental Mammal base wise conservation track by PhyloP.

**S2 Fig.**
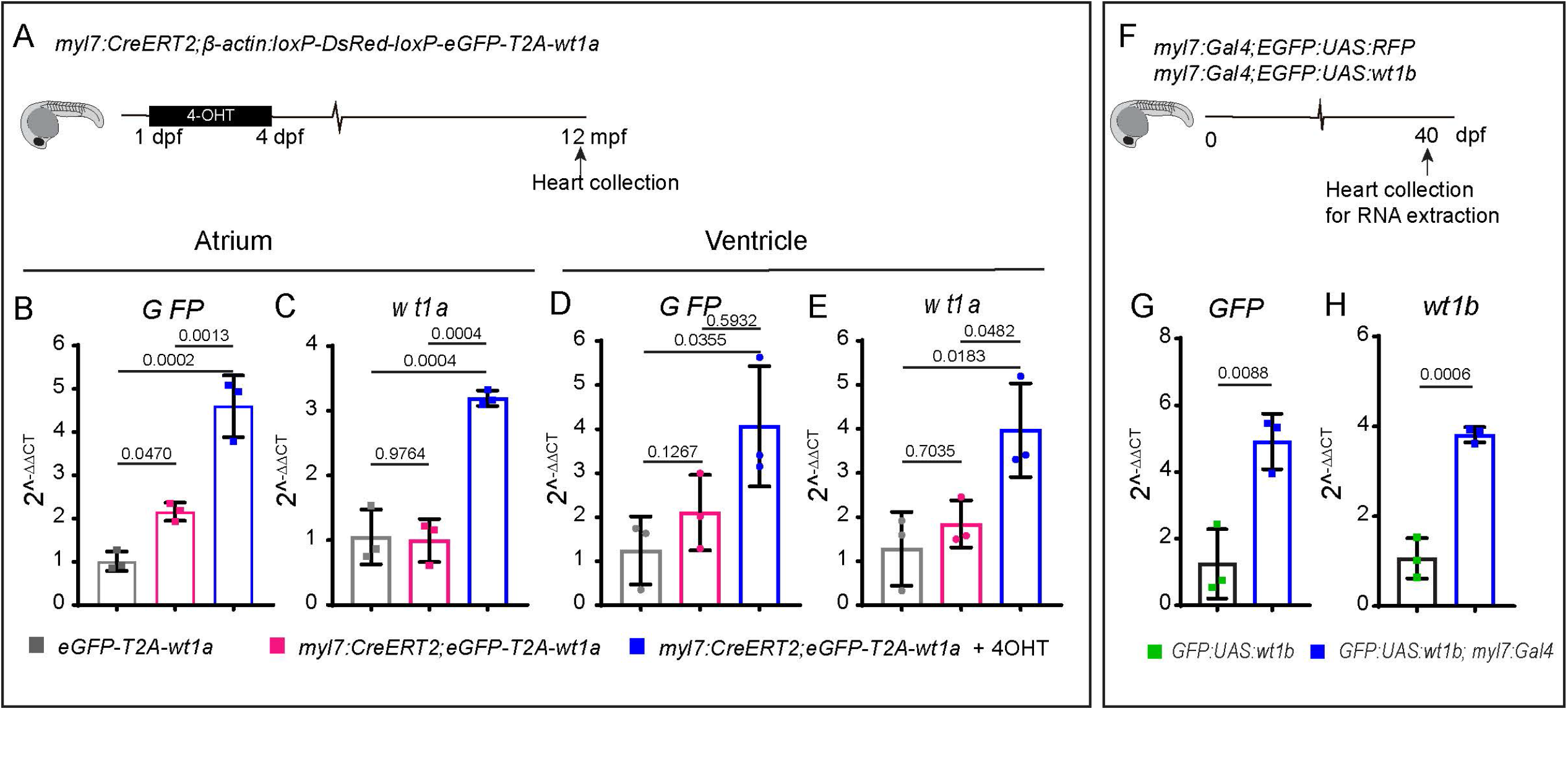
Validation of the *wt1a* and *wt1b* overexpression lines. (A) Schematic representation of the time points during which 4-Hydroxytamoxifen (4-OHT) was administered to *myl7:CreERT2;β-actin:loxP-DsRed-loxP-eGFP-T2A-wt1a* fish (in short *myl7:CreERT2,eGFP-T2A-wt1a*). (B-E) qRT-PCR for *GFP* (B,D) and *wt1a* (C,E) on adult heart cDNA from *myl7:CreERT2, eGFP-T2A-wt1a* with and without 4-OHT. qRT-PCR was performed on cDNA obtained from the atrium (B,C) and (D,E) ventricles. Points represent biological replicates, 3 for each group. Statistical significance was calculated using one-way ANOVA. Shown are means ±SD. (F) Schematic representation of lines used and the time at which RNA was extracted. (G-H) qRT-PCR for *eGFP* (G) and *wt1b* (H) in *eGFP:UAS:wt1b* and *myl7:Gal4;eGFP:UAS:wt1b* hearts at 40 days post fertilization (dpf). The points represent biological replicates. Statistical significance was calculated with an unpaired t-test. Shown are also means ±SD.

**S3 Fig.**
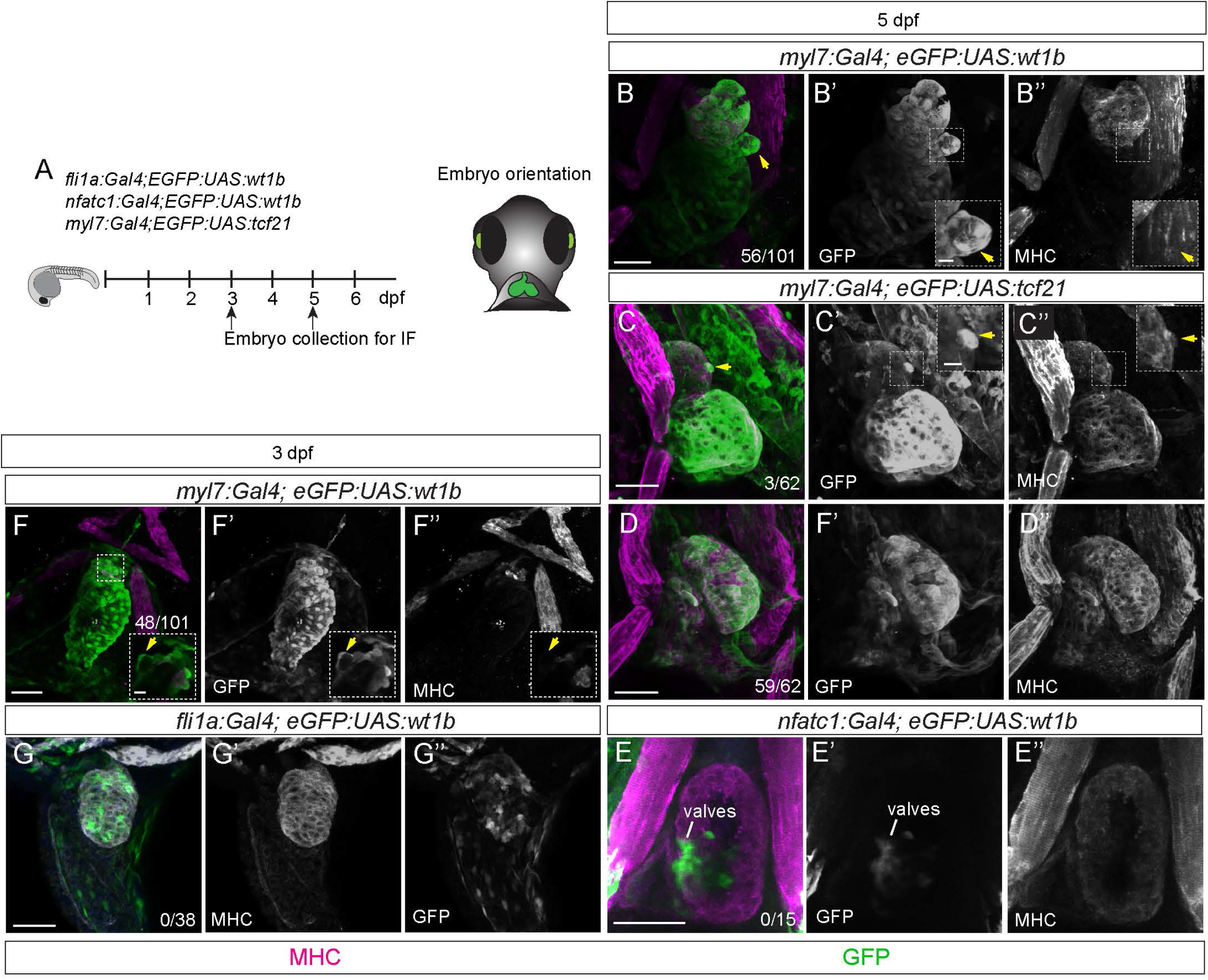
Overexpression of *tcf21* transcription factor in cardiomyocytes and *wt1b* in non-cardiomyocytes does not affect heart development. (A) Schematic representation of the lines used and embryo orientation for imaging. (B-G”) Immunofluorescence against GFP and myosin heavy chain (MHC) on *myl7:Gal4;eGFP:UAS:wt1b*, *myl7:Gal4;eGFP:UAS:tcf21, fli1a:Gal4;eGFP:UAS:wt1b* and *nfatc1:Gal4;eGFP:UAS:wt1b* zebrafish embryos, at 3 or 5 days post fertilization (dpf). (B-B”) Shown are 3D projections of a *myl7:Gal4;eGFP:UAS:wt1b* heart in a ventral view, at 5 dpf. (B’-B”) show single channels for GFP and MHC. The box highlights a zoomed region in the heart where a cluster of delaminating cells can be seen. (B”) Note the absence of MHC in the delaminated cells. (C-D”) Shown are 3D projections of a *myl7:Gal4;eGFP:UAS:tcf21* heart in a ventral view, at 5 dpf. (C-C”) show single channels for GFP and MHC. The box highlights a zoomed region in the heart with one cell delaminating. Note in C” that the delaminating cell preserved MHC expression. (E-E”) Shown are maximum intensity projections of 5 stacks with a distance of 1.5 µm between two consecutive optical sections of the heart region of a *nfatc1:Gal4;eGFP:UAS:wt1b* heart in a ventral view, at 5 dpf. GFP expression is observed in the valve region. The amount of embryos with delaminating cells is indicates in the panels. Green, GFP; magenta, MHC. Scale bar 50 µm and 10 µm, in the zoom boxes .at, atrium; v, ventricle.

**S4 Fig.**
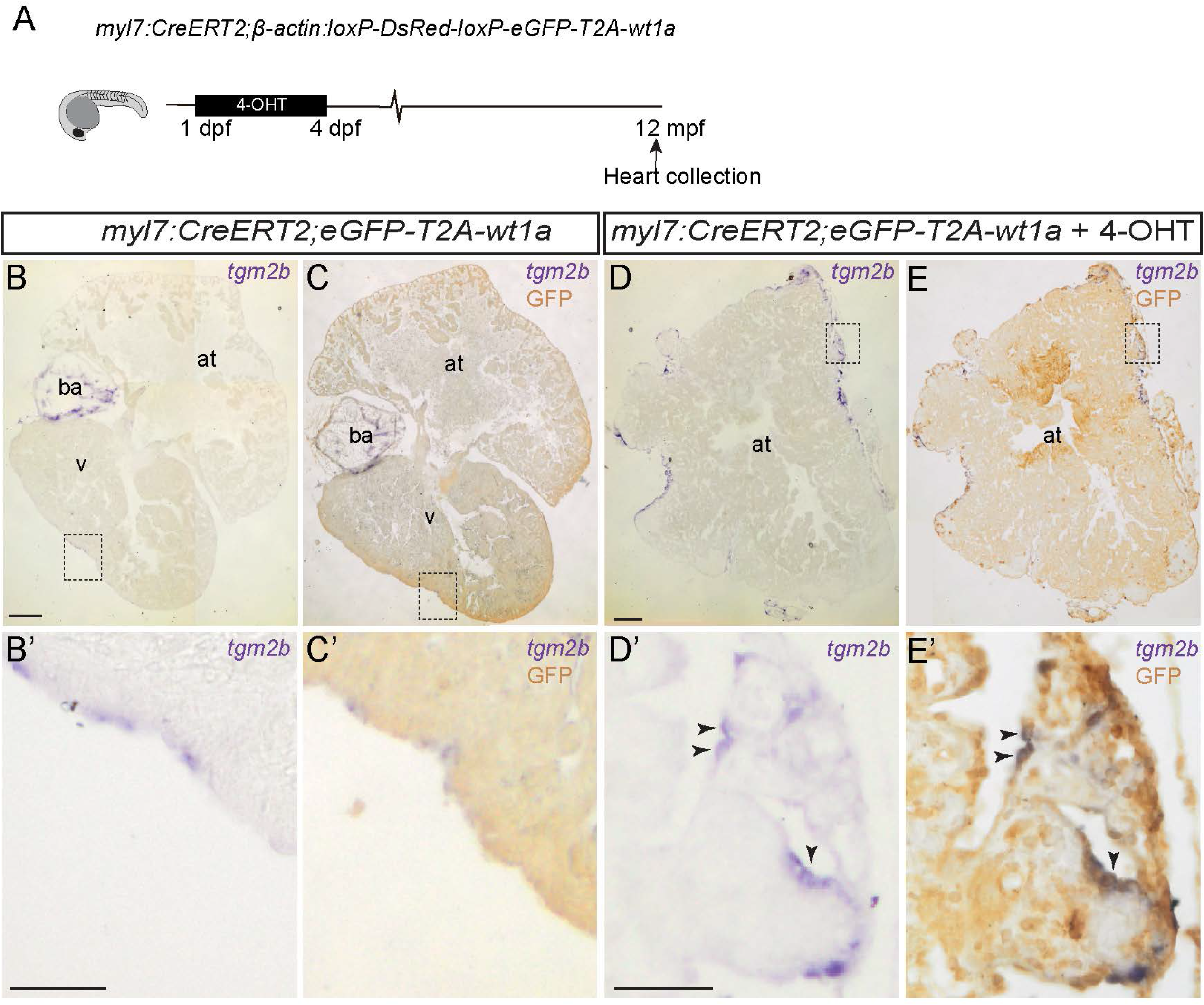
*wt1a* overexpression in cardiomyocytes express epicardial markers in the adult heart. (A) Schematic representation of the time points during which 4-hydroxytamoxifen (4-OHT) was administered to *myl7:CreERT2;β-actin:loxP-DsRed-loxP-eeGFP-T2A-wt1a* fish (in short *myl7:CreERT2,eeGFP-T2A-wt1a*). (B-E’) *in situ* mRNA hybridization against *tgm2b* and immunohistochemistry against eGFP on paraffin sections of *myl7:CreERT2,eeGFP-T2A-wt1a* (B-C’) and *myl7:CreERT2,eeGFP-T2A-wt1a* + 4-OHT (D-E’) adult hearts. (B,C,D and E) Images of sections after *in situ* mRNA hybridization against the epicardial marker *tgm2b.* (B’, C’, D’ and E’), same section as in B, C, D and E, after eGFP immunohistochemistry. Black arrows in E and E’ indicate double positive cells for *tgm2b* and eGFP. Scale bars: 200 µm (B,B’,D and D’), 50 µm (C,C’,E and E’).

**S5 Fig.**
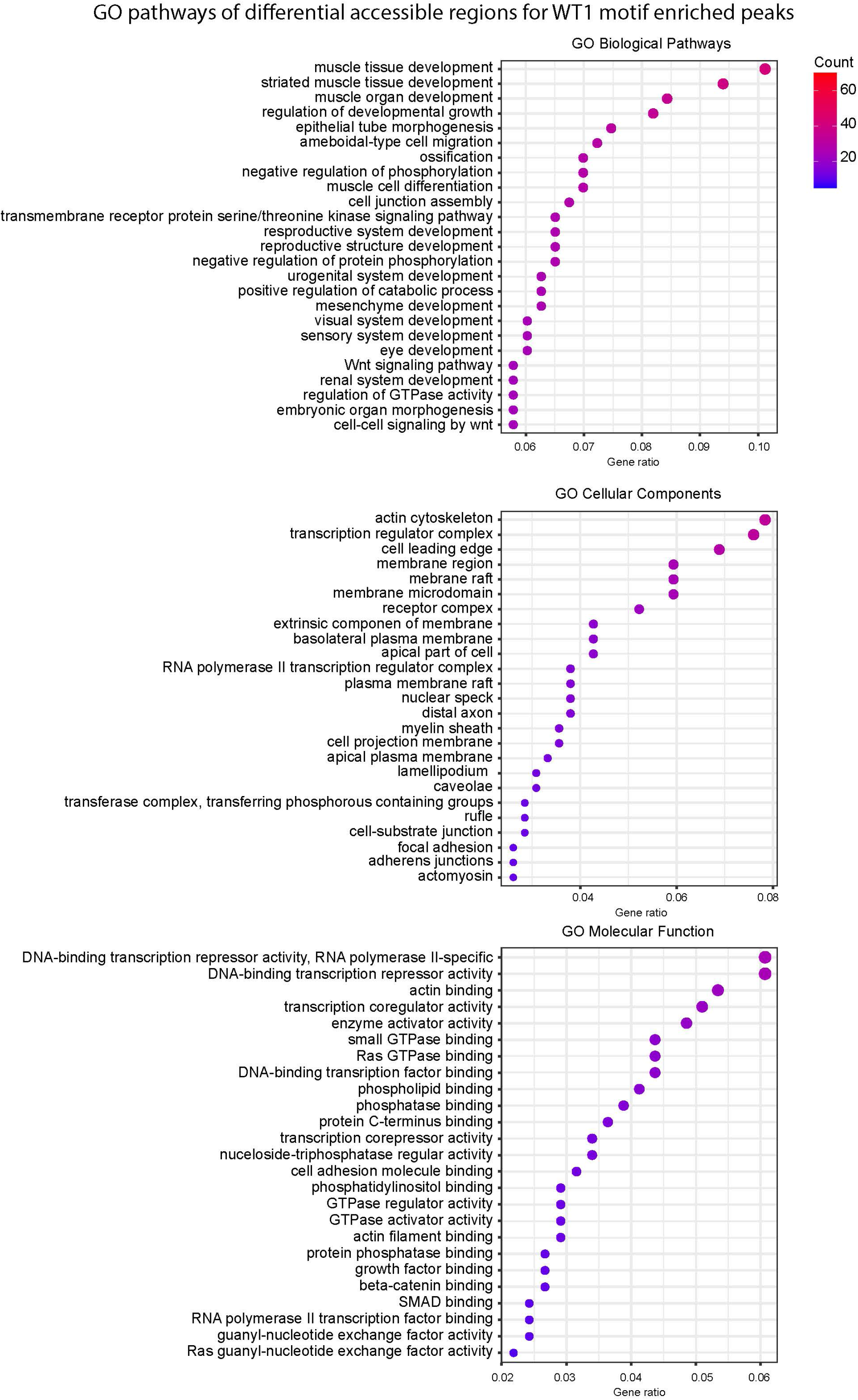
Gene Ontology pathways of differential accessible regions for WT1 motif enriched peaks. Gene Ontology (GO) pathways enrichment for differential accessible regions that contain the WT1 motif. Shown are the top 25 Biological, Cellular Components and the Molecular Function pathways.

**S6 Fig.**
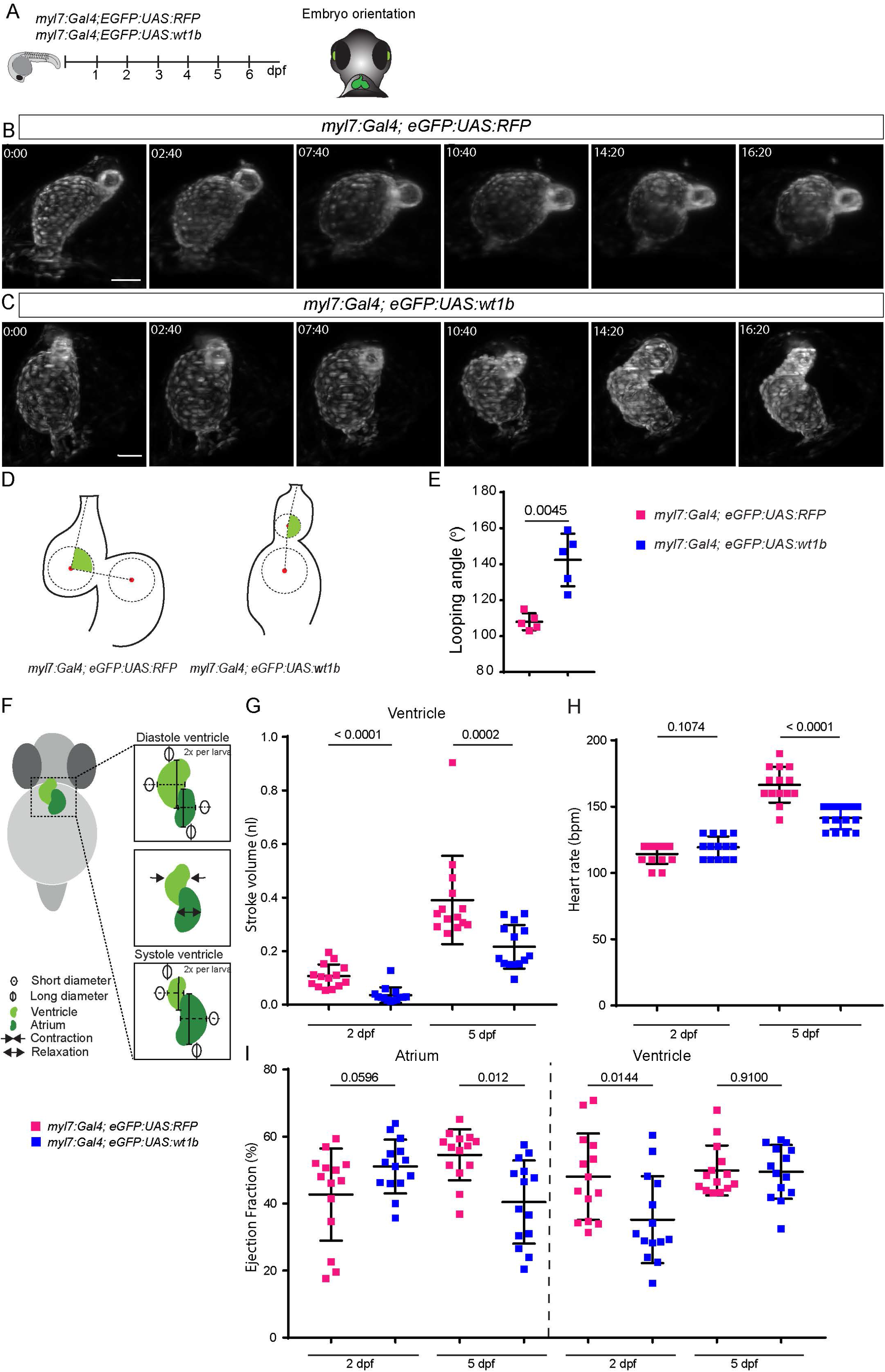
Heart looping and function are is impaired upon overexpression of wt1b in cardiomyocytes. (A) Schematic representation of the lines used and embryo positioning for image acquisition. (B-C) Time lapse images of heart looping in (B) *myl7:Gal4;eGFP:UAS:RFP* and (C) *myl7:Gal4; eGFP:UAS:wt1b* embryos between 2 and 3 days post-fertilization (dpf). Elapsed time since initial acquisition is stamped in each panel. Shown are ventral views with the head to the top. (D) Schematic representation of calculation of heart looping. (E) Quantification of the looping angle between the ventricle and the atrium at 5 dpf. Statistical significance calculated with unpaired t-test, with Welch’s correction. Each point represents one heart. Shown are means ±SD. (F) Schematic representation of parameters used to determine cardiac function in *myl7:Gal4;eGFP:UAS:RFP* and *myl7:Gal4; eGFP:UAS:wt1b*. (G) Quantification of ventricular stroke volume at 2 days post fertilization (dpf) and 5 dpf. Statistical significance was calculated with the Mann-Whitney test. (H) Quantification of the heart rate at 2 dpf and 5 dpf. Statistical significance calculated with an unpaired t-test. (I) Quantification of ventricular ejection fraction at 2 dpf and 5 dpf. Statistical significance was calculated with an unpaired t-test for the comparison between groups in the atrium and in the ventricle at 2 dpf. Mann-Whitney test was applied to calculate the statistical significance between the groups in the ventricle, at 5 dpf. In all graphs each point represents one embryo. Shown are also means ±SD. Scale bars, 50 µm. at, atrium; v, ventricle.

**S7 Fig.**
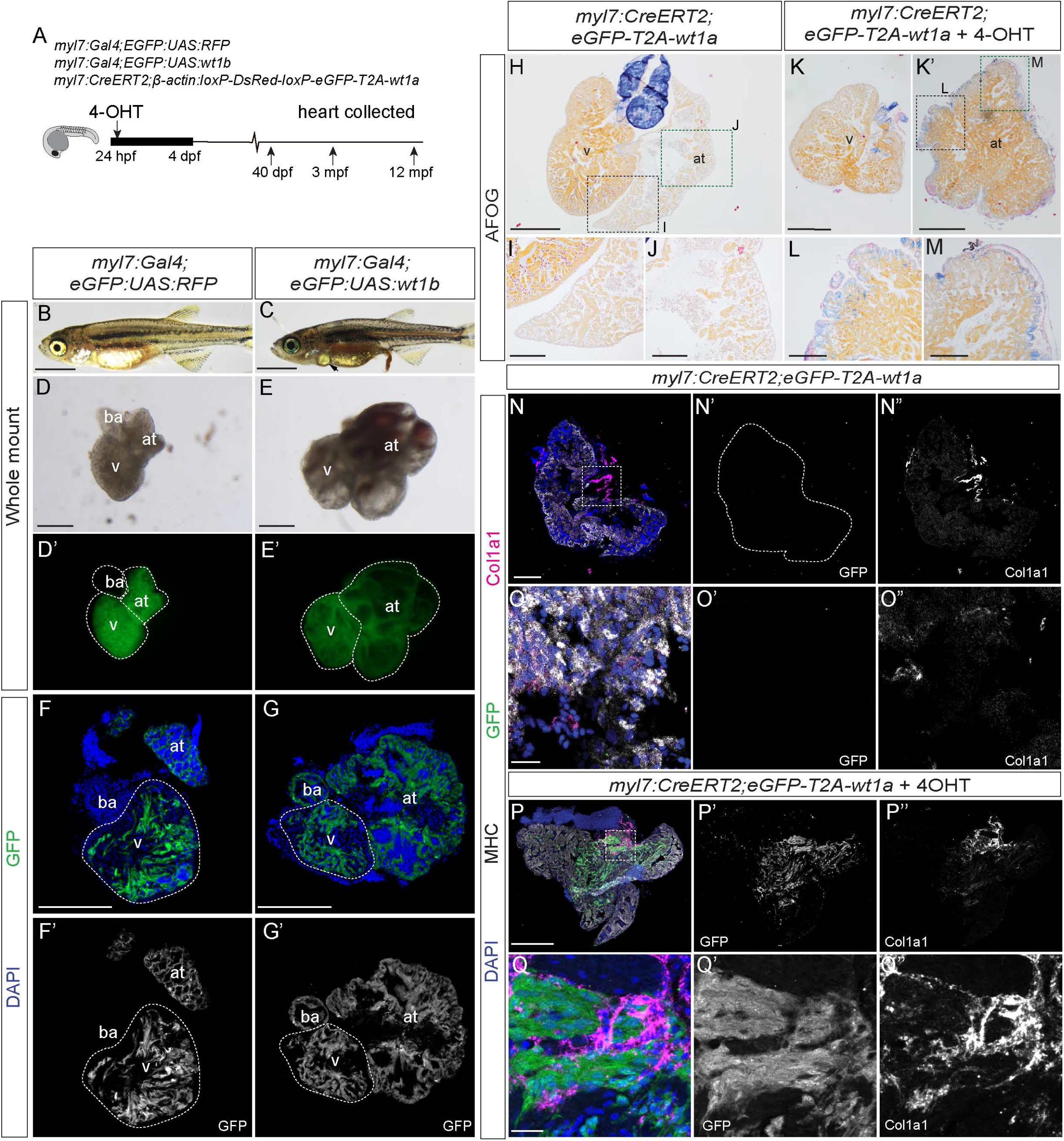
Morphological changes due to *wt1* overexpression in cardiomyocytes are sustained in larval and adult hearts. (A) Schematic representation of the lines used and the developmental stages at which hearts were analyzed. (B) 40 day post fertilization (dfp) juvenile *myl7:Gal4:eGFP:UAS:RFP* fish. (C) 40 dpf juvenile *myl7:Gal4:eGFP:UAS:wt1b* fish. Arrow points to pericardial edema. (D-D’) Dissected heart of a *myl7:Gal4:eGFP:UAS:RFP* and (E-E’) *myl7:Gal4:eGFP:UAS:wt1b* fish at 40 dpf. Note the enlarged and dysmorphic atrium. (F-F’) Midline section of a *myl7:Gal4:eGFP:UAS:RFP* and (G-G’) *myl7:Gal4:eGFP:UAS:wt1b* fish heart at 40 dpf. Note the high degree of myocardial tissue within the atrium of the *myl7:Gal4:eGFP:UAS:wt1b* heart. (H-M) AFOG staining on paraffin sections of *myl7:CreERT2,eGFP-T2A-wt1a* hearts non-recombined (H-J) or recombined (+ 4-OHT during embryogenesis) (K-M). (H) Whole heart section. (K) ventricle. (K’) corresponding atrium from the same animal. (I, J, L and M) Zoomed views of boxed areas in H and K. (N-Q”) Immunofluorescence against GFP (green), MHC (white) and Col1a1 (magenta) on adult atria cryosections of non-recombined (N-O”) and embryonically recombined (+4-OHT) *myl7:CreERT2,eGFP-T2A-wt1a* fish (P-Q”). (O-O”) Enlarged image of the boxed area in N. In *myl7:CreERT2,eGFP-T2A-wt1a* non recombined atria, Col1a1 staining is delimited to the valves and no GFP signal is detected. (Q-Q”) Enlarged image of the boxed area in P. In recombined *myl7:CreERT2,eGFP-T2A-wt1a* atria, Col1a1 staining is visible in myocardial areas close to GFP-positive cells. Scale bar: 1 mm (B and C); 500 µm (F-H and K); 200 µm (D-E’, I, J, L, M-N” and P-P”); 10 µm (O-O” and Q-Q”). at, atrium; v, ventricle; ba, bulbus arteriosus.

**S1 Video. *epi*:eGFP-positive cells at the venous pole switch off GFP expression and start expressing *myl7*:mRFP when entering the heart tube.**

*In vivo* time-lapse imaging of a *epi:GFP;myl7:mRFP* heart between 52 hpf and 68 hpf. The yellow arrow highlights a cell that initially is only GFP positive and latter stops expressing GFP and starts to express RFP. The cyan arrows point to cardiomyocytes in the heart tube that are still GFP -positive at the beginning of the video and then loose GFP signal concomitant with increase in mRFP signal intensity. Images were acquired with the Leica TCS SP8 DLS. Shown is a single plane reconstruction of the beating. Scale bar, 50 µm.

**S2 Video. Apical delamination of a *wt1b*-overexpressing cardiomyocyte in a cardiac ventricle at 2 dpf.**

*In vivo* time-lapse imaging of a *myl7:Gal4;eGFP:UAS:RFP* and a *myl7:Gal4;eGFP:UAS:wt1b* heart between 2 and 3 days post fertilization (dpf) acquired with the Leica TCS SP8 DLS confocal microscope, using the digital light sheet (DLS) mode. Shown is the reconstruction of a single plane of the beating ventricle. Note the rounded cells extruding from the ventricle in the *myl7:Gal4;eGFP:UAS:wt1b* heart (right panel, arrow). Scale bar, 50 µm.

**S3 Video. Apical delamination of a *wt1b*-overexpressing cardiomyocyte in a cardiac ventricle at 5 dpf.**

*In vivo* time-lapse imaging of a *myl7:Gal4;eGFP:UAS:wt1b* heart between 5 and 6 days post fertilization (dpf) acquired with the Leica TCS SP8 DLS confocal microscope, using the digital light sheet (DLS) mode. Shown is the reconstruction of a single plane of the beating ventricle. Note how the extruded cells flatten down during the time course of the video (Yellow arrow). Scale bar, 50 µm.

**S4 Video. Serial block face scanning z-stacks through a control zebrafish heart at 5 dpf.** Serial stacks through a ventricle from a *myl7:Gal4;eGFP:UAS:RFP* control embryo at 5 dpf. Images were obtained by serial block face scanning electron microscopy. Note the compact organization of the myocardium and the close connection between the myocardium and the endocardium, and how the sarcomeres form a continuous structure between adjacent cardiomyocytes. Of remark is also the dense border between the myocardium and epicardium. EpC, epicardial cell, EnC, endothelial cell, Ery, erythrocyte, CM, cardiomyocyte nuclei. Scale bar 10 µm.

**S5 Video. Zoomed view of a serial block face scanning z-stacks through a control zebrafish heart at 5 dpf.**

Serial stacks through a ventricle from a *myl7:Gal4;eGFP:UAS:RFP* control embryo at 5 dpf. Images were obtained by serial block face scanning electron microscopy. Shown is a magnification of the myocardium in a region where sarcomeres can be observed. Note the clearly marked z-lines and the longitudinal continuity of the sarcomeres between adjacent cardiomyocytes. Scale bar, 500 nm.

**S6 Video. Serial block face scanning z-stacks through a *wt1b*-overexpressing heart at 5 dpf.**

Serial stacks through a ventricle from a *myl7:Gal4;eGFP:UAS:wt1b* embryo at 5 dpf. Images were obtained by serial block face scanning electron microscopy. Note the absence of a compact and organized myocardial layer and the enlarged cardiac jelly separating the endocardium and the myocardium. Also visible the extensive areas filled with extracellular matrix. EpC, epicardial cell, EnC, endothelial cell, Ery, erythrocyte, CM, cardiomyocyte, ECM, extracellular matrix, v, ventricle, at, atrium. Scale bar, 20 µm.

**S7 Video. Zoomed view of a serial block face scanning z-stacks through a *wt1b*-overexpressing heart at 5 dpf.**

Serial stacks through a ventricle from a *myl7:Gal4;eGFP:UAS:wt1b* embryo at 5 dpf. Images were obtained by serial block face scanning electron microscopy. Shown here is a magnification of the myocardium in a region where sarcomeres can be observed. Note the z-lines. Remarkable is the disorganized arrangement of the sarcomeres between adjacent cardiomyocytes. Scale bar, 500 nm.

**S8 Video. Heart looping is impaired in *wt1b*-overexpression hearts.**

*In vivo* time-lapse imaging of a *myl7:Gal4;eGFP:UAS:RFP* and *myl7:Gal4;eGFP:UAS:wt1b* heart between 2 and 3 days post fertilization. Images were acquired with the Leica TCS SP8 DLS. Shown is a full 3D reconstruction of the beating heart through the looping process. Scale bar, 50 µm.

**S9 Video. z-stack through micro-computed tomography (µCT) acquisition of a *myl7:CreERT2;eGFP-T2A-wt1a* heart.**

Serial stack of a *myl7:CreERT2;eGFP-T2A-wt1a* heart obtained with a µCT scan of an adult thoracic cavity, used to evaluate the volume of the chambers of the heart. Marked are the bulbus arteriosus (ba), the ventricle (v) and the atrium (at). Scale bar, 500 µm.

**S10 Video. z-stack through micro-computed tomography (µCT) acquisition of a *myl7:CreERT2;eGFP-T2A-wt1a* heart recombined during embryogenesis.**

Serial stack of a *myl7:CreERT2;eGFP-T2A-wt1a* heart recombined between 24 hours post fertilization (hpf) and 4 days post fertilization (dpf), obtained with a µCT scan of an adult thorax, used to evaluate the volume of the chambers of the heart. Marked are the bulbus arteriosus (ba), the ventricle (v) and the atrium (at). Scale bar, 500 µm.

**S1 Data. Differential Peak Calling.**

The file contains about genomic location and information on fold change and significance values of the differential peaks. Columns J-O indicate in which samples peaks were identified (+) or not (-).

**S2 Data. Gene Onthology.**

Full list of pathways and genes enriched in each of them.

**S3 Data. Annotation of differential peaks.**

Differential peaks with their associated genes and genomic region classification.

**S1 Table.** Number of cells delaminating from the ventricle in the wt1a overexpression and control lines.

| Replicates | Number of delaminating cells | GFP+ | GFP- |
| --- | --- | --- | --- |
| Cross 1 | *myl7:CreErt2; wt1aOE-rec 1-4dpf-e1* | 11 | 0 |
|  | *myl7:CreErt2; wt1aOE-rec 1-4dpf-e2* | 13 | 0 |
|  | *myl7:CreErt2; wt1aOE-rec 1-4dpf-e3* | 17 | 0 |
|  | *myl7:CreErt2; wt1aOE-rec 1-4dpf-e4* | 12 | 0 |
|  | *myl7:CreErt2; wt1aOE-ctr-e1* | 0 | 0 |
|  | *myl7:CreErt2; wt1aOE-ctr-e2* | 0 | 0 |
|  | *myl7:CreErt2; wt1aOE-ctr-e3* | 0 | 0 |
|  | *myl7:CreErt2; wt1aOE-ctr-e4* | 0 | 0 |
|  | *myl7:CreErt2; wt1aOE-ctr-e5* | 0 | 0 |
|  | *myl7:CreErt2; wt1aOE-ctr-e6* | 0 | 0 |
| Cross 2 | *myl7:CreErt2; wt1aOE-rec 1-4dpf-e1* | 14 | 0 |
|  | *myl7:CreErt2; wt1aOE-rec 1-4dpf-e2* | 10 | 0 |
|  | *myl7:CreErt2; wt1aOE-rec 1-4dpf-e3* | 15 | 0 |
|  | *myl7:CreErt2; wt1aOE-rec 1-4dpf-e4* | 17 | 0 |
|  | *myl7:CreErt2; wt1aOE-rec 1-4dpf-e5* | 15 | 0 |
|  | *myl7:CreErt2; wt1aOE-rec 1-4dpf-e6* | 12 | 0 |
|  | *myl7:CreErt2; wt1aOE-ctr-e1* | 2 | 0 |
|  | *myl7:CreErt2; wt1aOE-ctr-e2* | 0 | 0 |
|  | *myl7:CreErt2; wt1aOE-ctr-e3* | 0 | 0 |
|  | *myl7:CreErt2; wt1aOE-ctr-e4* | 0 | 0 |

**S2 Table.** Number of animals with heart malformations in wt1b overexpression and control lines.

| Line Name | Replicates | GFP expression<br>*Tg/Tg* |  |  |  |  | No GFP expression<br>sibling |  |  |  |  |
| --- | --- | --- | --- | --- | --- | --- | --- | --- | --- | --- | --- |
|  |  | Normal Heart | Enlarged Atrium | Incorrect looping | Edema | Total | Normal Heart | Enlarged Atrium | Incorrect looping | Edema | Total |
| *myl7:Gal4; eGFP:UAS:wt1b* | Cross 1 | 26 | 18 | 18 | 18 | 44 | 135 | 0 | 0 | 2 | 137 |
|  | Cross 2 | 18 | 20 | 20 | 20 | 38 | 121 | 0 | 1 | 1 | 122 |
|  | Cross 3 | 2 | 24 | 26 | 26 | 28 | 97 | 0 | 0 | 1 | 98 |
|  | Cross 4 | 25 | 79 | 79 | 79 | 104 | 249 | 3 | 4 | 7 | 256 |
|  | Cross 5 | 6 | 41 | 37 | 38 | 47 | 146 | 0 | 0 | 6 | 152 |
|  | Cross 6 | 9 | 51 | 51 | 51 | 60 | 157 | 0 | 0 | 0 | 157 |
| *myl7:Gal4; eGFP:UAS:RFP* | Cross 1 | 8 | 1 | 1 | 1 | 9 | 53 | 0 | 0 | 1 | 54 |
|  | Cross 2 | 23 | 3 | 2 | 3 | 26 | 118 | 2 | 0 | 7 | 125 |
|  | Cross 3 | 10 | 4 | 2 | 4 | 14 | 46 | 0 | 2 | 4 | 50 |

**S3 Table.** Imaging settings: Parameters at Leica DLS for long term or high-resolution acquisition.

| Parameter | Long term imaging (Fig. ) |
| --- | --- |
| Objective lens detection/illumination | 25 x, NA 0.95 water immersion<br>2.5 x, NA 0.07 |
| Acquisition mode | XYTZL |
| Channel acquisition | Sequential, channels then stack |
| XY format | 512 x 512 (4 x 4 binning) |
| Exposure time, frame interval | 4.8 ms / 18.9 ms |
| T, number of frames | 50 |
| Z, optical slice interval | 2.5 μm |
| L, repeated acquisition | 99 Loops in 10 min intervals |
| Channel 1 laser intensity | 8% 488 nm |

